# Pleiotropic contribution of *rbfox1* to psychiatric and neurodevelopmental phenotypes in a zebrafish model

**DOI:** 10.1101/2023.02.23.529711

**Authors:** Ester Antón-Galindo, Maja Adel, Judit García-Gonzalez, Adele Leggieri, Laura López-Blanch, Manuel Irimia, William HJ Norton, Caroline H Brennan, Noèlia Fernàndez-Castillo, Bru Cormand

## Abstract

*RBFOX1* is a highly pleiotropic gene that contributes to several psychiatric and neurodevelopmental disorders. Both rare and common variants in *RBFOX1* have been associated with several psychiatric conditions, but the mechanisms underlying the pleiotropic effects of *RBFOX1* are not yet understood. Here we found that, in zebrafish, *rbfox1* is expressed in spinal cord, mid- and hindbrain during developmental stages. In adults, expression is restricted to specific areas of the brain, including telencephalic and diencephalic regions with an important role in receiving and processing sensory information and in directing behaviour. To investigate the effect of *rbfox1* deficiency on behaviour, we used *rbfox1*^sa15940^, a *rbfox1* loss-of-function line. We found that *rbfox1*^sa15940^ mutants present hyperactivity, thigmotaxis, decreased freezing behaviour and altered social behaviour. We repeated these behavioural tests in a second *rbfox1* loss-of-function line with a different genetic background, *rbfox1*^del19^, and found that *rbfox1* deficiency affects behaviour similarly in this line, although there were some differences. *rbfox1*^del19^ mutants present similar thigmotaxis, but stronger alterations in social behaviour and lower levels of hyperactivity than *rbfox1*^sa15940^ fish. Taken together, these results suggest that *rbfox1* deficiency leads to multiple behavioural changes in zebrafish that might be modulated by environmental, epigenetic and genetic background effects, and that resemble phenotypic alterations present in *Rbfox1*-deficient mice and in patients with different psychiatric conditions. Our study thus highlights the evolutionary conservation of *rbfox1* function in behaviour and paves the way to further investigate the mechanisms underlying *rbfox1* pleiotropy on the onset of neurodevelopmental and psychiatric disorders.

## INTRODUCTION

*RNA Binding Fox-1 Homolog 1* (*RBFOX1*, also referred to as *A2BP1* or *FOX1*) encodes an RNA splicing factor that is specifically expressed in brain, heart and muscle in human adults (GTEX, https://gtexportal.org/home/gene/RBFOX1). This gene regulates the expression and splicing of large gene networks and plays an important role in neurodevelopment ^1,2^. Rare genetic variations, including point mutations and copy number variants, have been reported in *RBFOX1* in patients with neurodevelopmental disorders such as autism spectrum disorder (ASD) ^3–6^, and *RBFOX1* haploinsufficiency results in a syndrome characterized by impaired neurodevelopment ^7,8^. In addition, transcriptomic analysis of brains from autistic individuals revealed decreased levels of *RBFOX1* and dysregulation of *RBFOX1*-dependent alternative splicing ^6,9^. *RBFOX1* has not only been related to neurodevelopmental conditions, but increasing evidence points to both rare and common variants in this gene as contributors to several psychiatric and neurological disorders ^5,6,10–12^. Interestingly, common variants in *RBFOX1* were found significantly associated with the cross-trait phenotype of the most recent genome-wide association studies (GWAS) meta-analysis of psychiatric disorders^13^ and *RBFOX1* was identified as the second most pleiotropic locus in a previous cross-disorder GWAS meta-analysis, showing association of common variants with attention-deficit/hyperactivity disorder (ADHD), autism spectrum disorder (ASD), bipolar disorder (BIP), major depression (MD), obsessive-compulsive disorder (OCD), schizophrenia (SCZ) and Tourette’s syndrome (TS) ^14^. Finally, *Rbfox1*^-/-^ mutant mice present a heightened susceptibility to seizures and neuronal hyperexcitability ^15^, and *Rbfox1* neuron-specific knockout mice show pronounced hyperactivity, stereotyped behaviour, impairments in fear acquisition and extinction, reduced social interest and lack of aggression ^6^, behaviours that are related to different psychiatric disorders. All these data suggest a major role for *RBFOX1* in psychopathology, although the mechanisms underlying its pleiotropic effects are not well understood.

In the last years, zebrafish have become a powerful model to study psychiatric disorders thanks to their high genetic similarity to human and their well-defined behavioural phenotypes, which can be easily assessed in the laboratory and compared to human psychiatric phenotypes ^16–18^. *RBFOX1* has two orthologous genes in zebrafish, *rbfox1* (*a2bp1*, NCBI gene ID: 449554) and *rbfox1l* (*a2bp1l*, NCBI gene ID: 407613). While the human gene *RBFOX1* is expressed in both the brain and skeletal and cardiac muscle (GTEX, https://gtexportal.org/home/gene/RBFOX1), *rbfox1* is mainly expressed in brain – but is also transcribed in heart –, and *rbfox1l* is exclusively expressed in skeletal and cardiac muscles at early developmental stages ^19,20^ and shows a low and restricted expression in only some neuronal populations of the adult zebrafish brain ^21^. In this study we focused on *rbfox1*, a gene that encodes a major protein isoform with 84% identity to the human protein ^19^, given its strong brain expression during development. To date, the expression of *rbfox1* at later stages has not been investigated nor its role in zebrafish neurodevelopment and behaviour.

Genetic studies in humans have pointed to a pleiotropic contribution of *RBFOX1* to several psychiatric conditions. Here, we have characterised the effect of loss of *rbfox1* function on zebrafish behaviour, and our data help describe mechanisms underlying its pleiotropic effects on the onset of neurodevelopmental and psychiatric disorders.

## MATERIAL AND METHODS

### Zebrafish strains, care and maintenance

Adult zebrafish and larvae (*Danio rerio*) were maintained at 28.5°C on a 14:10 light-dark cycle following standard protocols. All experimental procedures were approved by the Animal Welfare and Ethical Review board of the Generalitat de Catalunya. Behavioural experiments were performed using two different *rbfox1* mutant strains with different genetic backgrounds. *rbfox1*^sa15940^, on the Tübingen Long-fin (TL) background, is a transgenic line obtained from the European Zebrafish Resource Center of the Karlsruhe Institute of Technology (KIT). This line contains an intronic point mutation at the -2 position of a 3’ acceptor splicing site of *rbfox1* before the second/third exon of *rbfox1* annotated isoforms (A>T, Chr3:28068329, GRCz11) (Supplementary Figure 1). The second line, *rbfox1*^del19^, on the Tübingen (TU) background, was created using CRISPR/Cas9 genetic engineering and causes a frameshift deletion of 19 bp within exon 2 or 3 of *rbfox1* annotated isoforms (Chr3:28068264-28068282, GRCz11) (Supplementary Figure 2). Homozygous knockout fish (HOM, *rbfox1*^sa15940/sa15940^ and *rbfox1*^del19/del19^), heterozygous (HZ, *rbfox1*^sa15940/ +^ and *rbfox1*^del19/+^) and wild-type (WT, TL *rbfox1*^+/+^ and TU *rbfox1*^+/+^) fish were used for all behavioural experiments. For both *rbfox1*^sa15940^ and *rbfox1*^del19^ lines, homozygous, heterozygous and wild-type fish were obtained from heterozygous crosses to ensure a common genetic background.

### Gene expression analysis using Real-Time quantitative PCR (RT-qPCR)

Total RNA was extracted from the whole brain of 7 TL *rbfox1*^*+/+*^, 7 TL *rbfox1*^*sa15940/+*^ and 7 TL *rbfox1*^*sa15940 /sa15940*^ adult zebrafish and from 5 pools of 10 whole larvae of each genotype *rbfox1*^*+/+*^, *rbfox1*^*sa15940/+*^ and *rbfox1*^*sa15940 /sa15940*^ to perform RT-qPCR. Primers were designed to amplify cDNA from all the *rbfox1, rbfox1l, rbfox2, rbfox3a* and *rbfox3b* protein-coding isoforms described in the GRCz11 Ensembl database except for the *rbfox1*-203 isoform (http://www.ensembl.org/Danio_rerio/, Supplementary Table 1). In the case of *rbfox1* primers, the forward primer binds to the exon situated after the intronic point mutation in the *rbfox1*^*sa15940*^ line and the reverse primer binds to the subsequent exon. Results were normalised to the expression levels of reference housekeeping genes: the *ribosomal protein L13a* (*rpl13a*) and the *eukaryotic translation elongation factor 1 alpha 1a* (*eef1a1a*) housekeeping genes in the case of adult brains, and the *ubiquitin-conjugating enzyme E2A* (*ube2a*) in the case of larval samples, as the expression level of this gene is more stable through developmental stages (Supplementary Table 1). The relative expression of the genes and the fold change were calculated using the 2^-ΔΔCT^ comparative method ^22,23^.

#### *In situ* hybridization (ISH)

A specific mRNA probe targeting *rbfox1* (NCBI Reference Sequence: NM_001005596) was prepared and ISH experiments were performed in larvae (28 hours post fertilization (hpf), 2-, 3-, 4-, and 5-days post fertilization (dpf)) and dissected adult brains of WT fish from TL and TU lines. Further details are described in the Supplementary Methods.

### Generation of a *rbfox1* zebrafish loss-of-function line using CRISPR/Cas9

We used the CRISPR/Cas9 technology to generate stable *rbfox1* loss-of-function mutants (Supplementary Figure 2). Briefly, we designed 20 bp sequences (crRNA) targeting *rbfox1* next to a PAM sequence (Supplementary Table 2). scRNA, tracrRNA and Cas9 were purchased, and 1 nL of injection solution was injected into the cell of single-cell stage zebrafish embryos. After 24 hpf, the injection efficiency and crRNA efficacy were assessed and injected embryos (called F_0_ thereafter) with high injection efficiency were raised to adulthood. F_0_ were then crossed with WT zebrafish, generating F_1_ animals heterozygous for different mutations. DNA extraction and PCR followed by DNA Sanger sequencing analysis of F_1_ at 24 hpf identified the batches of F_1_ siblings that were more likely to contain a high proportion of frameshift mutations and the selected batches were raised to adulthood. F_1_ was screened to select a frameshift mutation and two siblings (one male and one female) that carried the same 19 bp frameshift mutation were in-crossed to generate F_2_ offspring that were 25% wild type, 50% heterozygous and 25% homozygous for the 19 bp mutation. The genotype of each F_2_ zebrafish was assessed to grow the animals and establish the mutant line. Further details of the method are described in the Supplementary Methods.

### Behavioural tests

A battery of behavioural tests was performed on adult zebrafish (3-6 months-old) using mixed groups of both sexes: open field test, shoaling test, visually-mediated social preference (VMSP) test, black and white test, and aggression test (Supplementary Figure 3). All the experiments were performed with homozygous knockout fish (*rbfox1*^sa15940/sa15940^ and *rbfox1*^del19/del19^), heterozygous (*rbfox1*^sa15940/+^ and *rbfox1*^del19/+^) and wild-type (TL *rbfox1*^+/+^ and TU *rbfox1*^+/+^) fish. In this first batch of experiments, the proportion of females in the tanks for both lines ranged from 50 to 70%. A second batch of experiments was performed with the *rbfox1*^sa15940^ fish segregating them by genotype and sex. Sex was phenotypically determined as it cannot be genotyped in zebrafish ^24,25^. In all instances, all fish were genotyped, sized-matched and 13 individuals were randomly selected per condition. These fish were then housed together in a single tank per condition for one week prior to and during the behavioural experiments. For each experiment, fish were placed in a different tank once tested to avoid retesting them and when the experiment was finished fish were housed together in the original tank. All tanks were kept in close proximity within the facility and were subjected to identical environmental and housing conditions. To prevent any potential housing bias, caretakers responsible for the fish were blind to both the genotype and experimental details.

Experiments were completed between 9:00 and 18:00 and recorded using StreamPix 7 software (Norpix) and a digital camera. Fish were left for 30 minutes to habituate to the testing room before the experiment. To determine the appropriate number of individuals in each group, we utilized GPower 3.1 software ^26^. The calculation was based on data from previous experiments conducted within the same experimental setup, aiming to guarantee sufficient statistical power to detect differences between the groups in the behavioural tests without using more animals than necessary. During the experiments, genotypes were alternated to prevent any bias resulting from the time of day or other possible confounders and all individuals were tested in the same setup and testing tanks. Most of the measures were performed automatically using a tracking system and analysed using software, ensuring blinding of the data. However, for a few measures that required manual quantification, a retrospective blinding process was implemented to ensure that there was no bias during the manual quantification of phenotypes. Further details of the tests are described in the Supplementary Methods.

### Statistical methods

Statistical analysis of RT-qPCR and behavioural data were performed with GraphPad Prism 8 (GraphPad Software, La Jolla California USA). The data sets were assessed for normality using D’Agostino-Pearson and Shaphiro-Wilk normality test and either a one-way ANOVA test followed by a Tukey’s post-hoc test or a Kruskal-Wallis test with Dunn’s correction for multiple testing were used to compare between multiple groups. Statistical analysis of the visually-mediated social preference test was performed by a two-way ANOVA with Sidak’s post-hoc test or a Kruskal-Wallis test followed by a Dunn’s correction for multiple testing. Standard deviation (SD) is indicated in the figures for each group of data. In the behavioural tests, the median of the individual speed was used instead of the mean as it was more representative of non-normal data caused by a high degree of freezing behaviour.

## RESULTS

### *rbfox1* expression is restricted to neurons during development and is localized to specific forebrain, midbrain and hindbrain areas in adulthood

During early development (28 hpf), *rbfox1* is expressed in spinal cord and hindbrain lateral neurons (Figure 1A). At later developmental stages (2-5 dpf) *rbfox1* expression is widespread in the mid- and hindbrain (Figure 1A). These findings are in line with previous published data ^27^. Furthermore, we found that during development *rbfox1* is also expressed in the heart, in line with what has previously been described elsewhere ^19^.

**Figure 1.**
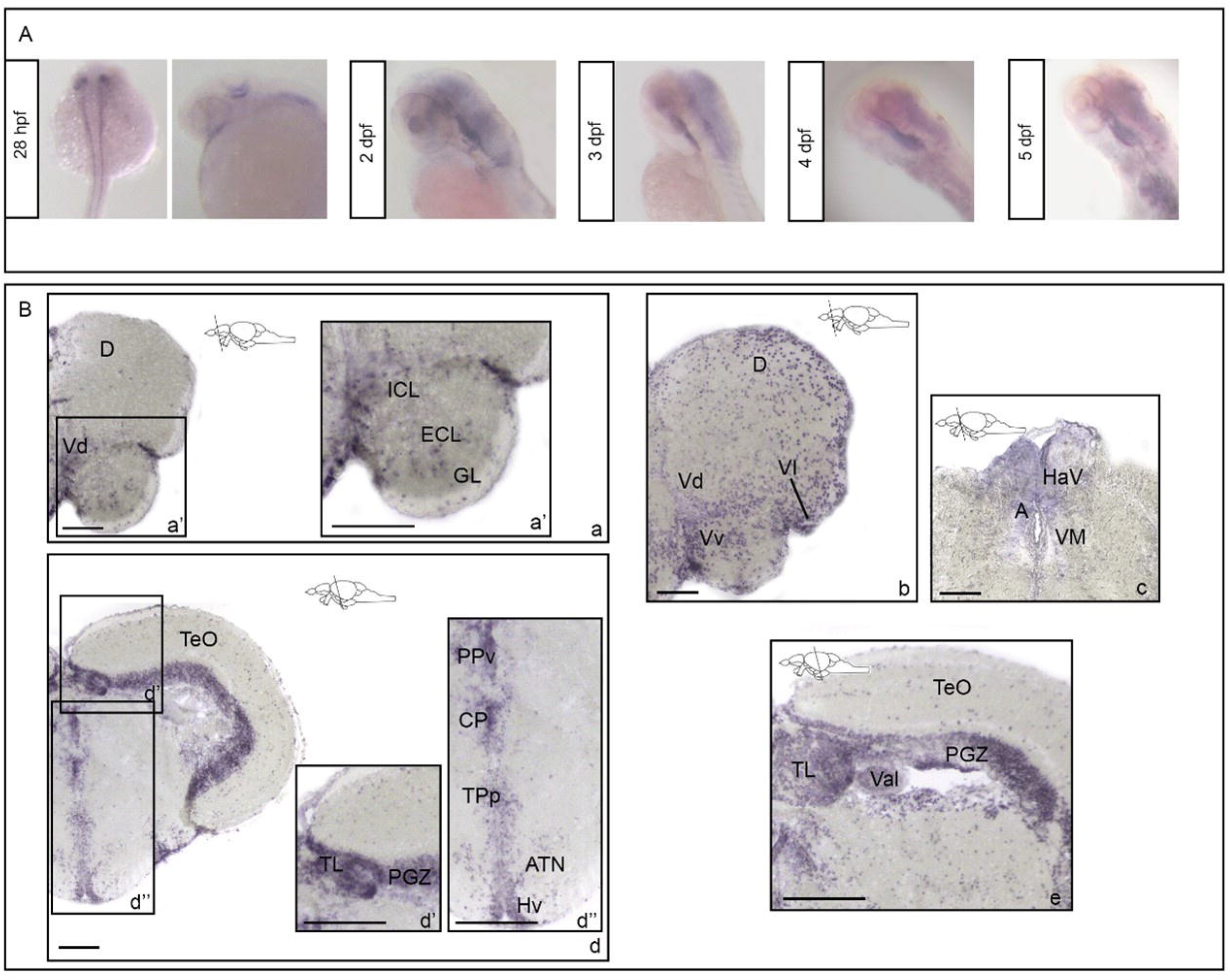
*rbfox1* shows restricted neuronal expression during development and is localized to specific forebrain, midbrain and hindbrain areas during adulthood. *rbfox1 in situ* hybridization on **A)** zebrafish whole mount larvae and **B)** adult zebrafish brains, TL background. **A)** *rbfox1* whole mount *in situ* hybridization on zebrafish larvae at 28 hours post fertilization, 3-, 4- and 5-days post fertilization. **B)** *rbfox1 in situ* hybridization on adult zebrafish brains, (a - c) forebrain and (d - e) midbrain transverse sections. A, anterior thalamic nuclei; ATN, anterior tuberal nucleus; CP, central posterior thalamic nucleus; D, dorsal telencephalic area; GL, glomerular cellular layer; HaV, ventral habenular nucleus; Hv, ventral zone of periventricular hypothalamus; ICL, internal cellular layer; PGZ, periventricular gray zone; PPv, ventral part of the periventricular pretectal nucleus; TeO, optic tectum; TL, torus longitudinalis; TPp, periventricular nucleus of posterior tuberculum; Val, valvula cerebelli; Vd, dorsal nucleus of ventral telencephalic area; Vl, lateral nucleus of ventral telencephalic area; VM, ventromedial thalamic nuclei; Vv, ventral nucleus of ventral telencephalic area. Scale bars: 100 μm (a, b, c, d); 200 μm (a’, d’, d’’, e).

In adult fish, *rbfox1* is expressed along the entire rostro-caudal brain axis. In the pallial region of the forebrain, *rbfox1* is expressed in the glomerular (GL), external (ECL) and internal (ICL) cellular layers of the olfactory bulbs (Figure 1B – a, a’). More caudally, *rbfox1* is expressed in the dorsal telencephalic area (D) and in the dorsal (Vd), lateral (Vl) and ventral (Vv) nuclei of ventral telencephalic area (Figure 1B – a, b). In the diencephalon, *rbfox1*-expressing cells have been detected in the ventral habenular nucleus (HaV), and in the anterior (A) and ventromedial (VM) thalamic nuclei (Figure 1B – c). *rbfox1* is also expressed in the periventricular layer of the thalamic and hypothalamic areas including the ventral part of the periventricular pretectal nucleus (PPv), the central posterior thalamic nucleus (CP), the periventricular nucleus of posterior tuberculum (TPp), the anterior tuberal nucleus (ATN), and the ventral zone of the periventricular hypothalamus (Hv) (Figure 1B – d, d’’). In the midbrain, *rbfox1* has been detected in the periventricular grey zone (PGZ) and in the torus longitudinalis (TL) (Figure 1B – d, d’, e). Finally, in the hindbrain *rbfox1* expression is observed in the lateral division of the valvula cerebelli (Val) (Figure 1B – e).

No differences were observed in *rbfox1* expression between TU and TL backgrounds (Supplementary Figure 4) at either larval or adult stages.

### *rbfox1*^*sa15940/sa15940*^ zebrafish do not express the complete *rbfox1* mRNA sequence and do not show alterations in the expression of the other *rbfox* genes

The first mutant line that we characterised, *rbfox1*^*sa15940*^ (A>T, Chr3:28068329, GRCz11), has an intronic point mutation at the -2 position of the 3’ acceptor splicing site before the second/third exon of all but one of the annotated *rbfox1* zebrafish isoforms (Supplementary Figure 1), which would cause that this exon is skipped during the splicing process and a shorter and aberrant *rbfox1* mRNA sequence produced. This mutation would cause the deletion of 81 aminoacids, which represent 22 to 54% of the original Rbfox1 protein isoforms, leading to a significant change in conformation that would affect the functionality of the mutant protein. By RT-qPCR, we observed a strongly decreased level of the expression of the *rbfox1* exon situated after the intronic point mutation in both homozygous and heterozygous *rbfox1*^*sa15940*^ mutant adult brains (93% and 43% respectively) compared to WT (mean HZ = 0.47; mean HOM = 0.07, WT vs. HOM: p = 0.0002, Figure 2A). We also observed a decreased level of the expression of this *rbfox1* exon in both homozygous and heterozygous *rbfox1*^*sa15940*^ mutant 3dpf larvae (80% and 38% respectively) compared to WT (mean HZ = 0.62; mean HOM = 0.20, WT vs. HOM: p = 0.0036, Figure 2D). These results suggest that this mutant line can be used to examine the effect of loss of *rbfox1* function in zebrafish.

**Figure 2.**
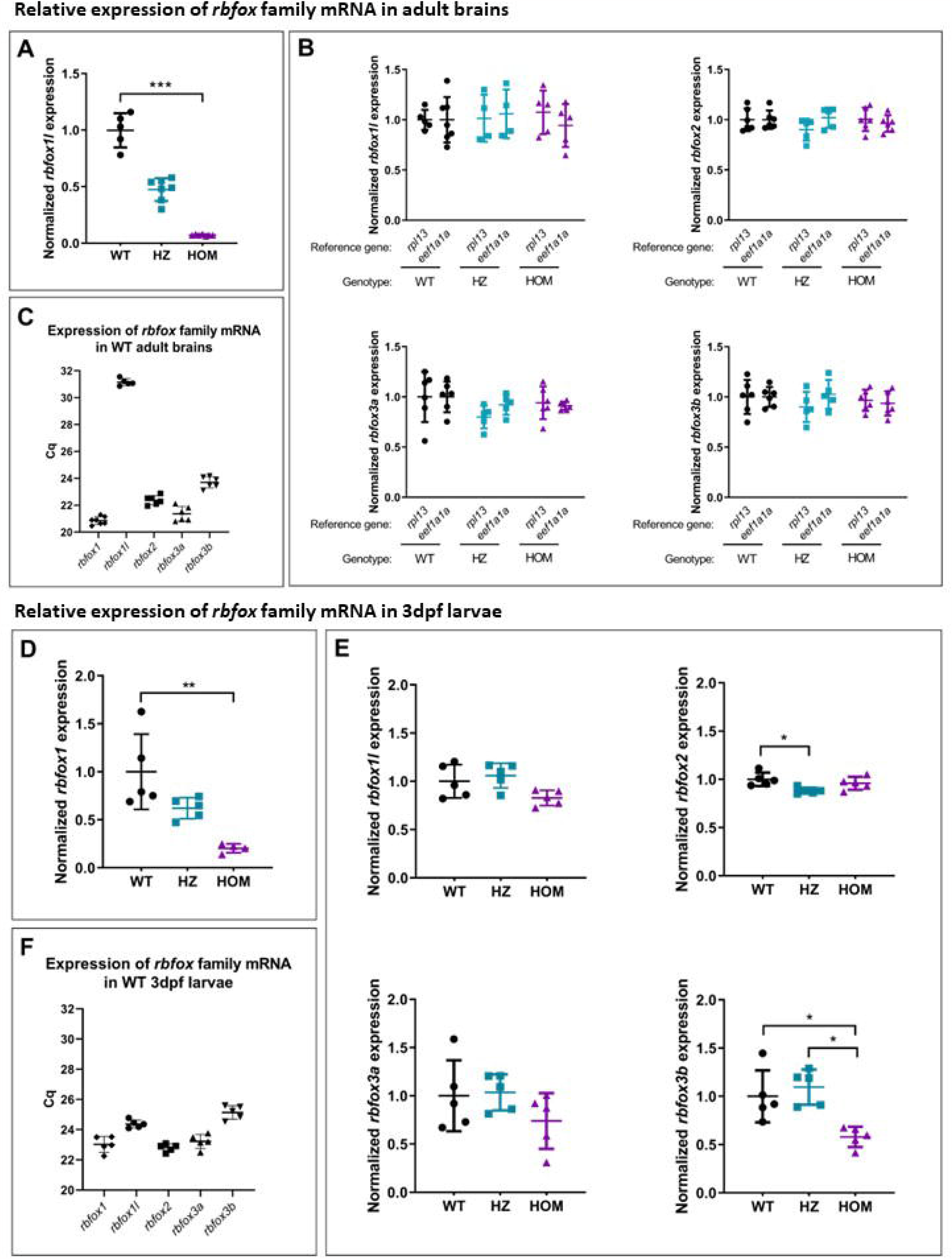
Relative expression of *rbfox* family mRNAs in adult brains and 3dpf larvae from the *rbfox1*^sa15940^ line. Adult brains: Relative brain expression of **A)** *rbfox1*, **B)** *rbfox1l, rbfox2, rbfox3a* and *rbfox3b* mRNA in adult brains. mRNA expression is normalised to the average expression of the mRNA in wild-type fish and to the reference housekeeping genes *ribosomal protein L13a* (*rpl13*) (A and B) or the *eukaryotic translation elongation factor 1 alpha 1a* (*eef1a1a*) (B). Kruskal-Wallis test followed by Dunn’s multiple comparison test. n = 5-7 WT, 5-7 HZ, 5-7 HOM. Mean ± SD. **C)** Brain expression of the *rbfox* family mRNAs in adult wild-type zebrafish brains represented as the quantification cycle value for each gene. n = 7 WT, 7 HZ, 7 HOM. Mean ± SD. **3dpf larvae:** Relative brain expression of **D)** *rbfox1*, **E)** *rbfox1l, rbfox2, rbfox3a* and *rbfox3b* mRNA in 3 dpf whole larvae. mRNA expression is normalised to the average expression of the mRNA in wild-type fish and to the reference housekeeping gene *ubiquitin-conjugating enzyme E2A* (*ube2a*). Kruskal-Wallis test followed by Dunn’s multiple comparison test. n = 5 WT, 5 HZ, 5 HOM. Mean ± SD. **F)** Brain expression of the *rbfox* family mRNAs in 3 dpf wild-type larvae represented as the quantification cycle value for each gene. n = 5 WT, 5 HZ, 5 HOM. Mean ± SD. Each point represents the results from a pool of 10 larvae.

We explored the expression levels of the four other *rbfox* zebrafish genes in both adult brains and 3dpf larvae. We observed no differences in the expression of *rbfox1l, rbfox2, rbfox3a* and *rbfox3b* in brain between WT and mutant *rbfox1*^*sa15940*^ adult fish (Figure 2B) and in the expression of *rbfox1l, rbfox2* and *rbfox3a* between WT and mutant *rbfox1*^*sa15940*^ larvae (Figure 2E). The expression of *rbfox3b* was slightly reduced in homozygous mutant *rbfox1*^*sa15940*^ larvae compared to WT and HZ (WT vs. HOM: p = 0.049, HZ vs. HOM: p = 0.011, Figure 2E). We also found that *rbfox1l* expression in the WT adult brain was very low compared to the expression of the other *rbfox* genes in this tissue, as its Cq (quantification cycle value) is much higher in the RT-qPCR analysis (Figure 2C). Unlike in adult brains, the expression levels of all the *rbfox* genes in whole WT 3dpf larvae were similar, with Cq ranging between 22 and 26 (Figure 2F).

Finally, the expression levels of the *rbfox* genes at 6dpf could not be assessed as all the reference genes considered (*rpl13a, eef1a1a, ube2a* and *tmem50a)* did not show stable expression among individuals and genotypes at this larval stage.

### Loss of *rbfox1* function produces behavioural alterations in *rbfox1*^*sa15940*^ zebrafish

We performed a battery of five behavioural tests (open field test, shoaling test, VMSP test, black and white test and aggression test) (Supplementary Figure 3) in TL WT *rbfox1*^*+/+*^, heterozygous (HZ) *rbfox1*^*sa15940/+*^, and homozygous (HOM) *rbfox1*^*sa15940/sa15940*^ adult fish, to investigate whether loss of *rbfox1* function affects behaviour (Figure 3 and Supplementary Figure 5).

**Figure 3.**
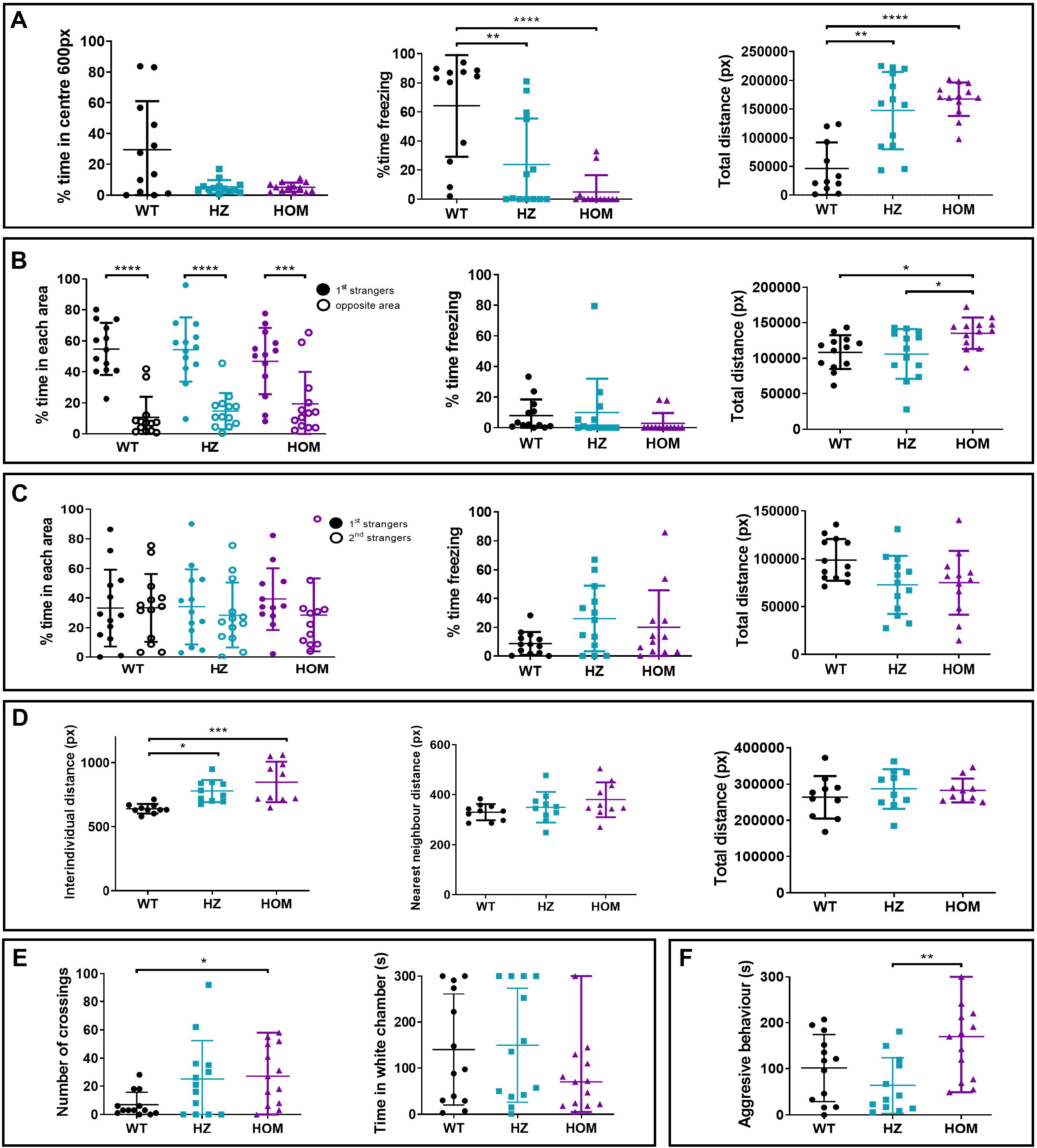
Behavioural alterations observed in the *rbfox1*^sa15940^ line. **A) Open field test**. Time spent in the centre of the arena, time spent freezing and total distance travelled during the open field test. One-way ANOVA followed by Tukey’s multiple comparison test. **B) Visually-mediated social preference test (VMSP). Social preference step**. Time spent in the area close to the 1^st^ strangers and in the opposite area, time spent freezing and total distance travelled during the social preference step of the VMSP test. Two-way ANOVA followed by Sidak’s multiple comparison test. **C) Visually-mediated social preference test. Preference for social novelty step**. Time spent in the areas close to the 1^st^ or 2^nd^ strangers, time spent freezing and total distance travelled during the preference for social novelty step of the VMSP test. Two-way ANOVA followed by Sidak’s multiple comparison test. **D) Shoaling test**. Mean of interindividual distance, nearest neighbour distance, cluster score and total distance travelled during the shoaling test. One-way ANOVA followed by Tukey’s multiple comparison test and Kruskal-Wallis followed by Dunn’s multiple comparisons test. **E) Black and white test**. Number of crossings between areas and time spent in the white area of the tank during the black and white test. Kruskal-Wallis followed by Dunn’s multiple comparisons test. **F) Mirror test**. Time spent exhibiting an aggressive behaviour against the mirror. For all the experiments except for the shoaling test: HOM, *rbfox1*^sa15940/sa15940^ fish; HZ, *rbfox1* ^sa15940/+^ fish; WT, wild-type TL. n = 13 WT, 13 HZ and 13 HOM for all tests except for the shoaling test. For the shoaling test: n = 2 groups of 5 individuals per genotype. * p < 0.05; ** p < 0.01; *** p < 0.001; **** p < 0.0001. Mean ± SD.

In this mutant line, all HZ and HOM individuals spend less than 20% of the time in the centre of the open field arena and show thigmotaxis, a behaviour that could be related to anxiety or stereotypies, whereas TL WT fish do not show preference to swim close to the walls of the arena (Figure 3A and Supplementary Figure 5A). In addition, HZ and HOM fish spend less time freezing than TL WT fish (WT vs. HZ, p = 0.0068; WT vs. HOM, p = 0.0001; Figure 3A) and show hyperactivity, as they swim longer distances (WT vs. HZ, p = 0.0027; WT vs. HOM, p = 0.0002; Figure 3A). They also present a higher swimming speed than TL WT individuals (WT vs. HZ, p = 0.0026; WT vs. HOM, p =0.0002; Supplementary Figure 5A).

In the visually-mediated social preference test (VMSP) we did not observe differences in social preference between genotypes for this line (Figure 3B and C, and Supplementary Figure 5B and C). In the first step, all the genotypes prefer to stay close to the group of stranger fish rather than in the opposite corner (1^st^ strangers vs. Opposite area: WT, p < 0.0001; HZ, p < 0.0001; HOM, p = 0.0005; Figure 3B) and in the second step all the genotypes show an equal preference for both stimulus groups (1^st^ strangers vs. 2^nd^ strangers: WT, p > 0.99; HZ, p = 0.90; HOM, p = 0.61; Figure 3C). However, in the first step of the test, mutant fish again showed hyperactivity, reflected by more distance travelled (HZ vs. HOM, p = 0.0282; WT vs. HOM, p = 0.0487; Figure 3B) and a higher speed of HOM fish compared to TL WT individuals (WT vs. HOM, p =0.0130; Supplementary Figure 5B).

In the shoaling test, we observed thigmotaxis in *rbfox1*^*sa15940*^ mutant fish (Supplementary Figure 5D) and we found differences in the mean interindividual distance (IID), which was higher in HZ and HOM compared to TL WT fish (WT vs. HZ, p = 0.0194; WT vs. HOM, p = 0.0005; Figure 3D). No differences were found in the time spent in the white chamber of the black and white test, but HOM fish cross more times the limit between areas, a sign of hyperactivity (WT vs. HOM, p = 0.0334; Figure 3E). Finally, no differences were observed between mutants and WT fish in the aggression test, but HOM fish are significantly more aggressive than HZ fish, as they spend more time exhibiting aggressive behaviour against a mirror (HZ vs. HOM, p = 0.0083; Figure 3F).

Taken together, these results show behavioural alterations in *rbfox1*^sa15940^ mutants in the TL genetic background that include hyperactivity, thigmotaxis and alterations in social behaviour.

We also investigated potential sex differences in the effects of *rbfox1*-deficiency using the *rbfox1*^sa15940^ line. In a second batch of experiments, fish were segregated by sex before and during the behavioural tests. We did not observe any significant variations in behaviour between males and females within the mutant groups, and only slight differences in locomotion during the initial step of the VMSP among the WT fish, where WT females travelled shorter distances and exhibited lower swimming speeds compared to WT males (Supplementary Figure 6). In this second batch, encompassing analyses of males, females, and combined groups, the mutant fish did not show hyperactivity or distinct thigmotaxis. However, alterations in social behaviour were evident in both VMSP and shoaling tests (Supplementary Figure 7A, B and C). Furthermore, we observed behavioural differences between the TL WT groups from the first and second batches (Supplementary Figure 7D), which could potentially account for the different outcomes obtained in these two batches..

### Loss of *rbfox1* function affects behaviour similarly in *rbfox1* ^*del19*^ fish

We then repeated the battery of behavioural tests in a second *rbfox1* mutant line with a TU genetic background, *rbfox1*^del19^, to investigate if *rbfox1*-deficiency affects behaviour also in this line. This line was created by using the CRISPR/Cas9 genome editing technique causing a frameshift deletion of 19 bp in exon 2 that disrupts the *rbfox1* coding sequence and produces a premature stop codon (Supplementary Figures 2 and 8). We observed behavioural differences between *rbfox1*^del19^ mutants and TU WT fish in all the tests performed, although some of the behavioural changes differed from those obtained for the *rbfox1*^sa19540^ line (Figure 4 and Supplementary Figure 9).

**Figure 4.**
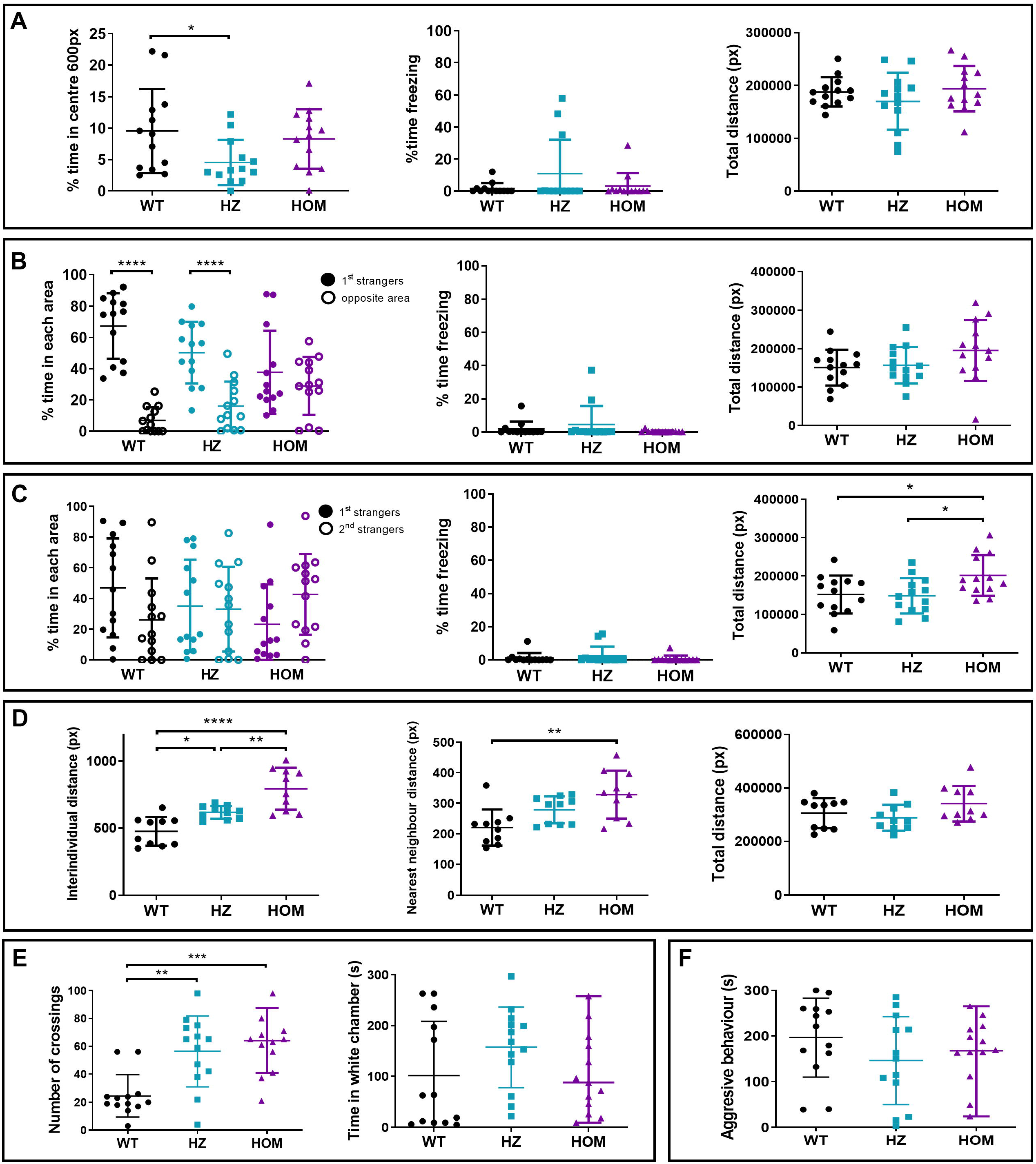
Behavioural alterations observed in the *rbfox1*^del19^ line. **A) Open field test**. Time spent in the centre of the arena, time spent freezing and total distance travelled during the open field test. One-way ANOVA followed by Tukey’s multiple comparison test. **B) Visually-mediated social preference test (VMSP). Social preference step**. Time spent in the area close to the 1^st^ strangers and in the opposite area, time spent freezing and total distance travelled during the social preference step of the VMSP test. Two-way ANOVA followed by Sidak’s multiple comparison test. **C) Visually-mediated social preference test. Preference for social novelty step**. Time spent in the areas close to the 1^st^ or 2^nd^ strangers, time spent freezing and total distance travelled during the preference for social novelty step of the VMSP test. Two-way ANOVA followed by Sidak’s multiple comparison test. **D) Shoaling test**. Mean of interindividual distance, nearest neighbour distance, cluster score and total distance travelled during the shoaling test. One-way ANOVA followed by Tukey’s multiple comparison test and Kruskal-Wallis followed by Dunn’s multiple comparisons test. **E) Black and white test**. Number of crossings between areas and time spent in the white area of the tank during the black and white test. Kruskal-Wallis followed by Dunn’s multiple comparisons test. **F) Mirror test**. Time spent exhibiting an aggressive behaviour against the mirror. For all the experiments except for the shoaling test: HOM, *rbfox1*^del19/ del19^ fish; HZ, *rbfox1*^del19/+^ fish; WT, wild-type TU. n = 13 WT, 13 HZ and 13 HOM for all tests except for the shoaling test. For the shoaling test: n = 2 groups of 5 individuals per genotype. * p < 0.05; ** p < 0.01; *** p < 0.001; **** p < 0.0001. Mean ± SD.

Similar to findings in *rbfox1* ^*sa15940*^, *rbfox1*^del19^ mutants tend to spend less time in the centre than TU WT fish, being significant for HZ fish (WT vs. HZ, p = 0.0467, Figure 4A) and present with thigmotaxis (Supplementary Figure 9A). However, we also observed differences in behaviour in the open field test between *rbfox1*^del19^ and *rbfox1*^sa15940^ lines: we did not find differences in locomotor activity (nor in distance travelled or speed) and freezing behaviour between genotypes in the *rbfox1*^del19^ line (Figure 4A and Supplementary Figure 9A).

In the preference step of the VMSP test, TU WT and HZ *rbfox1*^del19^ fish show a preference to stay close to stranger fish, whereas HOM *rbfox1*^del19^ fish show no social preference (1^st^ strangers vs. Opposite area: HOM, p = 0.6979; Figure 4B) and spend significantly less time than TU WT fish near strangers and more in the opposite area (WT vs. HOM, p =0.0057; Supplementary Figure 9B). In the social novelty preference step, we observed similar behaviour in both *rbfox1*^del19^ and *rbfox1*^sa15940^ lines: none of the genotypes show preference for a group of strangers (Figure 4C and Supplementary Figure 9C). In line with the *rbfox1*^sa15940^ results, HOM *rbfox1*^del19^ fish present hyperactivity in the two steps of the VMSP test, reflected by a higher speed (WT vs. HOM, p = 0.0339; Supplementary Figure 9B) and a further distance travelled (WT vs. HOM, p =0.0380; Figure 4C) than TU WT.

We found similar results in both *rbfox1* HOM lines in the shoaling and black and white tests: mutant *rbfox1*^del19^ fish present impaired social behaviour (IID: WT vs. HZ, p = 0.0235; WT vs. HOM, p < 0.0001; HZ vs. HOM, p = 0.0047; NND: WT vs. HOM, p < 0.0001; Figure 4D) and thigmotaxis (Supplementary Figure 9D) and HZ and HOM *rbfox1*^del19^ performed a higher number of crossings between areas than WT (WT vs. HZ, p = 0.0040; WT vs. HOM, p = 0.0006; Figure 4E).

Finally, in contrast to HOM *rbfox1*^sa15940^ fish, HOM *rbfox1*^del19^ fish were not more aggressive than HZ fish (Figure 4F).

In summary, both *rbfox1*^sa15940^ and *rbfox1*^del19^ mutants show hyperactivity, thigmotaxis and impaired social behaviour. However, each *rbfox1* line presents particularities: *rbfox1*^sa15940^ mutants show alterations in freezing behaviour and trends of aggression while *rbfox1*^del19^ mutants have stronger social impairments. The behavioural differences reported between the two *rbfox1* mutant lines might be explained by environmental effects and genetic background differences that modulate *rbfox1* effect on behaviour. Indeed, we can see that some behavioural aspects are different between the two WT lines, as we observe strong differences in the freezing behaviour (Supplementary Figure 10). Finally, even though discrepancies are reported, the effect of *rbfox1*-deficiency on behaviour in these two zebrafish models is in line with previous results found in *rbfox1*-deficient mice ^6^, as summarized in Table 1.

**Table 1:** Summary of behavioural alterations in *rbfox1*^sa15940^ and *rbfox1*^del19^ mutant zebrafish and *Rbfox1* depleted mice^6^

| Behavioural alterations | Zebrafish TL (♀, ♂) | Zebrafish TU (♀, ♂) | Mice (♂) |
| --- | --- | --- | --- |
| **Hyperactivity** | Total distance and swimming speed increased in HZ and HOM fish in the Open Field test<br>Total distance and swimming speed increased in HOM fish in the VMSP test<br>Number of crossings increased in HOM fish in the Black and White test | Total distance and swimming speed increased in HOM fish in the VMSP test<br>Number of crossings increased in HZ and HOM fish in the Black and White test | Total distance increased in HOM mice in the Open field test with and without the presence of a novel object in the centre<br>Total distance increased in HOM mice in the Light-dark box test<br>Total distance increased in HOM mice in the Marble-burying test |
| **Increased thigmotaxis behaviour** | Thigmotaxis increased in HZ and HOM fish in the shoaling and Open Field test | Thigmotaxis increased in HZ fish in the Open Field test<br>Thigmotaxis increased in HZ and HOM fish in the Shoaling test | Thigmotaxis increased in HOM mice in the Open Field test<br>Thigmotaxis decreased in HOM mice in the Novel Object Exploration paradigm |
| **Decreased freezing behaviour** | Decreased freezing in HZ and HOM fish in the Open Field | No differences found between the genotypes | Decreased freezing in HOM mice in the fear conditioning test in all stages<br>Decreased freezing in HZ mice in the fear conditioning test in extinction phase |
| **Decreased social interest** | Increased interindividual distance in HZ and HOM fish in the Shoaling test | Increased interindividual distance in HZ and HOM fish in the Shoaling test<br>Increased nearest neighbour distance in HOM fish in the Shoaling test<br>HOM fish spend less time near the 1<sup>st</sup> strangers in first step of the VMSP paradigm | Less social interest in social interaction test in the HOM mice |
| **Altered aggressive behaviour** | No difference between mutant and control fish in aggressive behaviour in the mirror test, but aggression is increased in HOM fish compared to HZ fish | No differences found between the genotypes | Lack of aggressive behaviour in HOM mice in the escalated aggression test |

HOM, homozygous individuals for *rbfox1* loss-of-function (LOF) mutation; HZ, heterozygous individuals for *rbfox1* loss-of-function (LOF) mutation; VMSP, visually-mediated social preference test; WT, wild-type individuals: ♀, female individuals; ♂, male individuals.

## DISCUSSION

In this study we have investigated the role of *rbfox1* in neurodevelopmental and psychiatric disorders by studying the behavioural effects of loss of *rbfox1* function in zebrafish. This gene has previously been reported to be highly pleiotropic, contributing to several psychiatric disorders ^13,14,28^. In addition, we have validated zebrafish *rbfox1*^sa15940^ and *rbfox1*^del19^ HOM lines as models of neurodevelopmental and psychiatric conditions.

First, *rbfox1* shows a restricted expression in brain and heart across developmental stages that suggests an important role of this gene during brain zebrafish development, in line with previous findings. Indeed, a study in human neural progenitor cells demonstrated that *RBFOX1* regulates splicing and expression of large gene networks implicated in neuronal development and maturation ^29^, and another study showed that *Rbfox1* controls synaptic transmission in the mouse brain ^15,30^. Also, previous studies in mice have shown that specific *Rbfox1* deficiency in the central nervous system leads to impairments in neuronal migration, axon extension, dendritic arborisation and synapse network formation, suggesting that loss of *Rbfox1* function contributes to the pathophysiology of neurodevelopmental disorders ^31–33^. Finally, several point mutations and copy number variations (CNVs) in *RBFOX1* have been described in patients with neurodevelopmental disorders, such as ASD and ADHD ^4–6,10,34^. We therefore hypothesise that loss of *rbfox1* function may affect brain maturation in zebrafish and therefore lead to impaired neuronal function and transmission during adulthood, with implications in the sensory response to the environment and in behaviour.

In addition, *rbfox1* specific expression is found mainly in forebrain areas in adult WT zebrafish, including the dorsal and ventral telencephalon, thalamus and periventricular hypothalamus. Interestingly, these areas are involved in receiving and processing sensory information, stress, and in directing behaviour, especially social behaviour and emotion ^35–38^. Given the important role of Rbfox1 in controlling splicing and expression in neurons, *rbfox1* deficiency may induce an impaired neuronal function in these areas with an impact on sensory processing, stress and behaviour in zebrafish.

Interestingly, both *rbfox1*^sa15940^ and *rbfox1*^del19^ HOM lines present alterations in behaviour. *rbfox1*^sa15940^ mutants present hyperactivity, thigmotaxis – a behaviour related to anxiety and stereotypies –, decreased freezing behaviour and altered social behaviour. *rbfox1*^del19^ mutants present similar thigmotaxis, but stronger alterations in social behaviour and lower levels of hyperactivity than *rbfox1*^sa15940^ fish. These results are in line with the behavioural alterations observed in a neuron-specific *Rbfox1* KO mouse line that presents decreased *Rbfox1* expression, as *Rbfox1* KO mice show a pronounced hyperactivity, thigmotaxis and reduced social interest ^6^. All these behavioural phenotypes can be assimilated to phenotypic alterations observed in patients with psychiatric or neurodevelopmental conditions. For example, social impairment is a symptom of ASD, hyperactivity of ADHD, and thigmotaxis is considered an anxiety-like behaviour in mouse and zebrafish.

We found differences in behaviour between *rbfox1*^sa15940^ and *rbfox1*^del19^ lines. On one hand, *rbfox1*^sa15940^ is a hyperactive line that presents with thigmotaxis and slight social impairments. On the other hand, *rbfox1*^del19^ fish present also with thigmotaxis, show only hyperactivity in one of the tests performed, and present stronger social impairments than *rbfox1*^sa15940^ fish. The phenotypic variations observed between the two zebrafish lines are likely attributable to genetic background disparities and/or environmental influences, although we cannot rule out differential effects of the two *rbfox1* mutations (e.g. exon skipping in the *rbfox1*^sa15940^ line versus a frameshift deletion in the *rbfox1*^del19^line). Another factor that may contribute to the behavioural differences found between lines is the tank effect: each genotype was segregated in a separate tank and therefore behaviour could be influenced by the conspecifics present in the tank. In addition, behavioural differences between WT TL and TU strains have been previously reported, WT TL fish being considered more anxious and sensitive to anxiogenic stimuli than TU WT fish^39^. Our results are in line with these reported phenotypes, as we found that TL WT presents a strong freezing behaviour, especially in the open field test, that is not present in TU WT fish. In addition, *Rbfox1* KO mice present behavioural alterations not described in the zebrafish lines such as, lack of aggressive behaviour, and behaviours that could not be tested in our zebrafish lines for practical reasons such as deficit in the acoustic startle response and impairments in fear acquisition ^6^. Given the differences observed between the WT zebrafish lines, we hypothesised that loss of *rbfox1* function alters behaviour differently depending on other environmental and genetic effects.

Moreover, when separating *rbfox1*^sa15940^ fish by sex in a second batch of experiments, the results obtained were different from the first batch, although social behaviour was shown to be altered as well. Indeed, the WT fish from the two batches behave differently in some tests, being more active in the second batch. These differences might be explained again by the influence of the environment and the genetic background. The fish used in this second batch come from a new generation of *rbfox1*^sa15940^ fish that were bred with a different TL WT strain and it is known that zebrafish strains are not completely inbred and genetically well-defined as it is the case with laboratory mice ^40^, which might lead to variations in the genetic background between these two batches. In addition, housing fish in sex-separated groups before and during the experiments has been described to affect behaviour ^41^.

These results suggest that, on one side, environmental effects might play a role when assessing behavioural effects of a genetic variation and, on the other side, that the effects of variants in other genes may contribute to the final phenotype, in agreement with a recent proposed genetic model for complex psychiatric disorders composed by ‘hub’ and ‘peripheral’ genes ^42–45^. Our results show that the damaging effect of a loss-of-function mutation in *rbfox1* may be modulated by genetic and environmental effects and therefore lead to different phenotypes, which is also in line with the different diagnosis of patients with rare CNVs or point mutations in the *RBFOX1* gene as well as the contribution of common variants to different psychiatric disorders ^6,10,11,14,46^.

To conclude, all these results show that loss-of-function of *rbfox1* in zebrafish and mice leads to behavioural alterations that can be related to different neurodevelopmental and psychiatric disorders. Thus, our data contribute to a better understanding of the involvement of *RBFOX1* in psychiatric disorders and point to a pleiotropic contribution of this gene that can be modulated by other environmental and genetic factors. In addition, we have validated two new *rbfox1* HOM zebrafish lines to be used as models for psychiatric disorders, in which further experiments can be performed to unravel the molecular mechanisms that link *RBFOX1* with psychiatric phenotypes.

## AUTHORS CONTRIBUTION

N.F-C. and B.C. conceived and coordinated the study. N.F-C. and B.C. designed the experimental approaches for the behavioural experiments. E.A-G. designed and conducted the behavioural experiments, contributed to the characterization of the loss-of-function lines and wrote the paper. M.A. conducted the second batch of behavioural experiments and the larvae qPCR experiments. J.G-G. designed and performed the CRISPR/Cas9 experiment. A.L. designed and conducted the ISH experiments and contributed to the characterization of the loss-of-function lines. L.L-B. and M.I. contributed to the behavioural experiments. W.H.J.N contributed to the design of the behavioural experiments. CH.B. supervised the CRISPR/Cas9, ISH and qPCR experiments. All authors discussed and commented on the manuscript.

## ACKNOWLEDGEMENTS

*rbfox1*^sa15940^ zebrafish embryos were generated and obtained from the European Zebrafish Resource Center of the Karlsruhe Institute of Technology (KIT). Major financial support for this research was received by BC from the Spanish ‘Ministerio de Ciencia, Innovación y Universidades’ (RTI2018-100968-B-100, PID2021-1277760B-I100), the ‘Ministerio de Sanidad, Servicios Sociales e Igualdad/Plan Nacional Sobre Drogas’ (PNSD-2017I050 and PNSD-2020I042), ‘Generalitat de Catalunya/AGAUR’ (2021-SGR-01093), ICREA Academia 2021, and the European Union H2020 Program [H2020/2014-2020] under grant agreements n° 667302 (CoCA) and Eat2beNICE (728018), and received by CHB from the NIH (USA) (U01 DA044400-03). E.A-G was supported by the Ministerio de Economía y Competitividad (Spanish Government), the EU H2020 program (Eat2beNICE-728018) and a Margarita Salas postdoctoral grant. M.A. was supported by the ‘Studienstiftung des Deutschen Volkes’. J.G-G. was supported by the Queen Mary Principal’s Research Studentship in the School of Biological and Chemical Sciences.

## CONFLICT OF INTEREST

The authors declare no conflict of interest.

**Supplementary information is available at TP’s website**.

