## Supplementary Material for "Pleiotropic contribution of *rbfox1* to psychiatric and neurodevelopmental phenotypes in a zebrafish model"

1. **SUPPLEMENTARY METHODS**
2. **SUPPLEMENTARY FIGURES**
3. **SUPPLEMENTARY TABLES**
4. **REFERENCES**
1. **SUPPLEMENTARY METHODS**

**In situ hybridization**

*In situ hybridization on whole mount zebrafish larvae*

*In situ* hybridization was carried out on 28 hours post fertilization (hpf), 2- 3- 4- and 5-days post fertilization (dpf) larvae of the TU wild-type strain. To prevent skin pigmentation, embryos were incubated in 0.2 mM 1-phenyl 2-thiourea (PTU) (Sigma, Gillingham, UK) from 24 hpf. When they reached the desired ages, larvae were fixed in 4% paraformaldehyde (PFA) (Sigma, Gillingham, UK) to avoid tissue degradation, overnight (ON) at 4°C. The following day, larvae were rinsed in 1x phosphate buffered saline (PBS) supplemented with Tween (0.05% v/v), dehydrated in an ascending methanol series (25%, 50%, 70%, 80%, 90%, 100% methanol, 5 min each), and stored in 100% methanol at -20 °C. To perform *in situ* hybridization experiments, larvae were rehydrated in descending methanol series (100%, 90%, 80%, 70%, 50%, 25% methanol), 5 min each, and washed in 1x PBS, 5 min. After washes, larvae were permeabilized using proteinase K (stock 20 μg/mL in PBS) as follows: 28 hpf larvae were permeabilized in 1x proteinase K for 20 min at room temperature (RT), older stages were permeabilized with 2x proteinase K at 37°C for 30 min. Then, larvae were post fixed in 4% PFA for 20 min and washed in 1x PBS at RT, 5 x 5 min. Prehybridization was carried out in a hybridization solution (HB) containing 50% formamide, 5% saline sodium citrate buffer (SSC), 50 µg/mL heparin, 0.05 mg/mL yeast RNA, 0.1% Tween 20, and 0.92% citric acid at 68 °C for 2 h. Thereafter, sections were incubated in HB containing *rbfox1* riboprobe, ON at 68°C. Post hybridization washes were performed at 68°C with a gradient of 2x SSC and formamide (75%, 50%, 25% and 0% formamide), 10 min each, and then twice with 0.02x SSC, 30 min each. Subsequently, larvae were blocked in blocking solution (BS) containing 10% normal sheep serum and 2 µg/µL bovine serum albumin, for 1 h at RT. After blocking step, larvae were incubated in a 1:2000 dilution of anti-digoxigenin Fab fragments conjugated with alkaline phosphatase (Roche) in BS, 1h at RT and then ON at 4°C. The following day, larvae were washed in 1x PBS, 6 x 15 min each, and then in NTMT (100 mM Trish HCl, 50 mM MgCl_2_, 100 mM NaCl, 0.1% Tween), 3 x 5 min each. The chromogenic reaction was carried out by incubating the larvae in NBT/BCIP solution (Sigma-Aldrich by Merck) in NTMT buffer, at RT in the dark, and were observed every 20 min until the signal detection. After the signal was developed, larvae were washed in 1x PBS at RT, post fixed in 4% PFA for 20 min, cleared and stored in 80% glycerol at 4°C. Pictures were acquired by Leica MZ75 microscope.

*In situ hybridization on adult brain sections*

*In situ* hybridization on adult zebrafish was conducted on paraffin embedded wild type brains, in the TU and TLF backgrounds. Fish were culled by overdose of tricaine prior to head removal. Brains were dissected and fixed in 4% PFA (Sigma, Gillingham, UK) in 1x PBS, ON at 4°C. Brains were then rinsed in 1x PBS and dehydrated in ascending ethanol series (15 min in each of 30%, 50%, 70%, 80%, 90%, 100% ethanol) and embedded in paraffin. Transverse sections of 12 µm thickness were cut using a microtome (Leica, Wetzlar, DE). To perform *in situ* hybridization, slides were de-waxed in xylene (twice, 10 min each), rehydrated in descending ethanol series (2 x 5 min in absolute ethanol, then 90%, 80%, and 70% ethanol, 5 min each), and rinsed in 1x PBS for 5 min. After washes, sections were permeabilized using proteinase K (0.05 μg/μL), for 8 min at RT. Proteinase K action was then inactivated by two washes in 2 mg/mL glycine, 5 min each. Sections were post fixed in 4% PFA for 20 min and washed in 1x PBS at RT. Prehybridization was carried out in HB, for 1h at 68°C. Thereafter, sections were incubated in HB containing *rbfox1* riboprobe, ON at 68°C. Post hybridization washes were performed at 68°C twice for 20 min in 1x SSC, twice for 20 min in 0.2x SSC, and several washes were performed in 1x PBS, 5 min each at RT. Then, sections were blocked in BS for 30 min at RT and incubated in a 1:2000 dilution of anti-digoxigenin Fab fragments conjugated with alkaline phosphatase (Roche) in BS, ON at 4°C. The following day, sections were washed in 1x PBS, 5 x 10 min each. The chromogenic reaction was carried out by incubating the slides in NBT/BCIP solution (Sigma-Aldrich by Merck) in NTMT, at RT in the dark, and were observed every 20 min until the signal detection. After the signal was developed, sections were washed in 1x PBS at RT, dehydrated in ascending ethanol series (70%, 80%, 90%, 100% ethanol, 5 min each), cleared in xylene (twice, 5 min each) and mounted with mounting medium. Pictures were acquired using a Leica DMRA2 upright epifluorescent microscope with colour QIClick camera (Leica, Wetzlar, Germany) and processed with Velocity 6.3.1 software (Quorum Technologies Inc). Anatomical structures were identified according to the Neuroanatomy of the Zebrafish Brain by Wullimann et al., 1996 ^1^.

**Generation of a *rbfox1* zebrafish loss-of-function line using CRISPR/Cas9**

*Design of crRNA targeting genes of interest*

The genome browser Ensemble (www.ensembl.org) was used to find suitable crRNAs sites targeting early coding exons. All the crRNAs were created in regions with overlapping sites for restriction enzymes. This enabled us to detect when Cas9 cut the DNA and imperfect DNA repair happened, so that the target site for the restriction enzyme was lost. The crRNAs designed to target the DNA regions of interest are included in **Supplementary Table 2**.

Any restriction enzyme can be used as long as it covers one nucleotide on either side of the CRISPR cut size. However, we selected MwoI (recognition sequence -GCNNNNNNNGC-, New England Biolabs, Cat.R0573S) whose recognition sequence includes a PAM site (NGG or CCN) and covered wide regions. This enzymes function in standard PCR reactions without using special buffers.

*Preparation of crRNA, tracrRNA and Cas9 protein*

crRNAs and tracrRNA (Sigma, Cat.TRACRRNA05N) were purchased. Upon arrival, crRNAs and tracrRNA were re-suspended in nuclease free water at a concentration of 250 ng/μL and separated into 5 μL aliquotes using RNAse free PCR tubes. Each tube was not freeze thawed more than three times and they were stored at -20°C for short term use and -80°C for long term storage. We also bought Cas9 protein (New England Biolabs, Cat.M0386M) that was diluted 1:3 in buffer B (New England Biolabs, Cat.B802S) on arrival, and stored at -20°C.

*Embryo injection*

1 nL of injection solution was injected into one-cell stage zebrafish embryos. We assembled a 4 μL total injection solution as follows:

| Injection mix: |  |
| --- | --- |
| crRNA (250ng/μL) | 1 μL |
| tracrRNA (250ng/ μL) | 1 μL |
| Cas9 protein (1:4 from stock) | 1 μL |
| Phenol red (1:10) | 1 μL |

100 to 150 embryos were injected. From those, eight were used to assess the consistency of the injection technique and crRNA efficacy. The remaining embryos were raised. Un-injected sibling controls were not raised to adulthood, but they were kept as controls for 24 hours.

*Assessment of injection and crRNA efficacy*

We collected eight injected embryos and eight un-injected sibling controls at 24 hpf (we waited for 24 hours because the CRISPR/Cas9 system can still create double strand breaks hours after the injection). Embryos were individually placed in 96-well PCR plates, all the water was removed, and DNA was extracted using the Hot Shock procedure and PCR amplification with Biomix Red (Bioline, Cat.BIO-25006). Primer sequences used for PCR genotyping were: forward, 5’-CCAGGCAGTCATTTGACCAT-3’; reverse 5’-CAAGTTCAGCGTGTGCTCTG-3’. Once the PCR was completed, 1 μL of restriction enzyme (diluted 1:4 in H_2_O) was added to the PCR reaction of four uninjected control embryos (to confirm that the digestion was successful) and eight injected embryos (to evaluate the injection technique and crRNA efficacy). We then incubated the reaction at the optimal cutting temperature for the enzyme.

A high number of injected zebrafish carrying mutations increases the probability of finding large genomic lesions in the F_1_. More than four out of eight injected zebrafish presented incomplete digestion, suggesting that Cas9 had cut the target region and therefore the restriction site was lost. We injected ∼100 embryos and raised ∼50 to cover for unexpected variables such as high death rates during raising.

*F_0_ screen*

Injected F_0_ animals were ready after 3-4 months of age to test whether frame-shift mutations were transmitted to the offspring. F_0_ zebrafish are usually mosaic animals, which means that there are mixed genotypes (mutant and wild type) in the same injected animal. Since the gametes generated by F_0_ zebrafish can also contain different genotypes, a F_0_ screening was carried out to assess the frequency and type of mutations transmitted to the offspring. The F_0_ screen consisted of a cross of a F_0_ zebrafish with a wild type, generating F_1_ animals heterozygous for different mutations.

∼20 F_0_ zebrafish were individually crossed with wild type zebrafish. For each F_0_ pair-wise outcross with wild-type, we collected 50 to 100 embryos and from those, we took a random sample of eight embryos to extract their DNA at 24 hpf and amplify by PCR the region of interest.

We confirmed by PCR that F_0_ zebrafish from one pair transmitted relatively large mutations to the offspring. Two or three out of the eight F_1_ embryos screened contained amplicons with ∼20 bp difference in size from the wild-type amplicon (detected by a band shift on the agarose gels). We kept these F_0_ progenitors isolated until the transmitted mutations were confirmed.

Due to the mosaic nature of the injected F_0_, the germ cells may transmit different mutations in each gamete and give rise to different F_1_ mutants. Some F_1_ zebrafish may carry frameshift mutations whereas other F_1_ zebrafish may carry mutations that do not alter the amino acid sequence. To ensure that the F_1_ do carry frameshift mutations, we 1) cloned the PCR products of interest to isolate the mutant amplicons, 2) transformed competent cells with the cloning product and incubated them ON at 37 °C, 3) selected the positive colonies and confirmed the insertion of the amplicon in the TOPO vector by PCR using M13 forward: 5’-GTAAAACGACGGCCAG-3’ and reverse 5’-CAGGAAACAGCTATGAC-3’ primers and 4) sent the candidate amplicons to be Sanger sequenced by Source BioScience PLC.

The sequencing results confirmed that these embryos carried a 19 bp frameshift deletion for *rbfox1* (**Supplementary Figure 1, panel B**). ∼50 F_1_ embryos, offspring of a F_0_ by wild type pair, were raised to adulthood. We assumed that, if two/three embryos drawn from a sample of n=8 had the same mutation, it would be likely to find at least two sibling zebrafish (ideally a male-female pair) with the same frameshift mutation in n∼50.

*F_1_ screen, F_2_ genotyping and breeding of F_3_*

When F_1_ embryos reached maturity, we performed a F_1_ screen to select the alleles that would establish the loss-of-function line (LoF). We fin clipped F_1_ zebrafish, heterozygous for different mutations, to extract their DNA. We then genotyped them by PCR, ran the product on a gel to identify those carrying large mutations, cloned and sequenced the PCR products of interest to confirm that the insertions/deletions were causing a sequence frameshift. The procedure for PCR, cloning, transformation, colony PCR, and sequencing was the same than for the F_0_ screen. We identified one male and one female with 19 bp frameshift mutations for *rbfox1* (**Supplementary Figure 1, panel C**). Since these were heterozygous animals, we incrossed them to obtain F_2_ offspring that would be 25% homozygous, 50% heterozygous and 25% wild type for LoF *rbfox1*. Before conducting behavioural assays, we outcrossed and genotyped by PCR the mutant lines for at least two further generations to remove potential off-target mutations.

**Behavioural tests**

A battery of five behavioural tests was performed on adult zebrafish (**Supplementary Figure 2**):

*Open field test*

The open field test was performed in a large circular open tank (43 cm diameter) and the fish were recorded from above for 5 minutes. We used idtracker.ai ^2^ and the *trajectorytools* module for Python (<https://github.com/fjhheras/trajectorytools>) to quantify the time spent in the centre of the tank (i.e. the inner 50% of the area), the time spent freezing, the distance swum and the velocity. We used 13 individuals per genotype.

*Shoaling test*

The shoaling experiment was performed following the protocol from Parker et al., 2013 ^3^. We used idtracker.ai and the *trajectorytools* module for Python to measure the nearest neighbour distance (NDD), the inter-individual distance (IID), the distance swum and the velocity. We also virtually divided the tank into nine sections and calculated the cluster score across time ^3^. We used two groups of five individuals per genotype.

*Visually-mediated social preference test (VMSP)*

The experiment was performed in a transparent tank composed of one central chamber (20 cm x 14 cm) surrounded by four identical chambers (10 cm x 7 cm). This test is divided in two steps as described in Carreño Gutierrez et al., 2019 ^4^: social preference step and preference for social novelty step. During the first step (social preference), a first group of three unfamiliar wild-type fish were placed into one of the side compartments. Then, the behaviour of a focal fish placed in the central area was recorded for five minutes. The time spent closer to the first group of strangers was compared to the time spent near the empty area diagonally opposite. In a second step (social novelty preference), a second group of three unfamiliar zebrafish were placed in the compartment diagonally opposite the first group. The focal fish was recorded for 5 more minutes. We used a mixture of size-matched males and females as strangers since they can attract both male and female zebrafish ^5^. We used idtracker.ai and the *trajectorytools* module for Python to measure the time spent in the different areas of the central compartment, the time spent freezing, the distance swum and the velocity. We used 13 individuals per genotype.

*Black and white test (scototaxis test)*

The black and white test was performed in a rectangular tank (24 cm x 12 cm) divided into two equal areas, a black area and a white area. Fish were placed in the centre of the tank and recorded for 5 minutes. The time spent in each area and the number of crossings between them were manually quantified. We used 13 individuals per genotype.

*Aggression test*

Aggression was measured using the mirror-induced stimulation protocol ^6^. A single fish was placed in the centre of tank with three white walls and a transparent wall, through which an external mirror can be seen, and was recorded for 5 minutes. The time spent in agonistic interaction with the mirror was manually quantified. We used 13 individuals per genotype.

2. **SUPPLEMENTARY FIGURES**

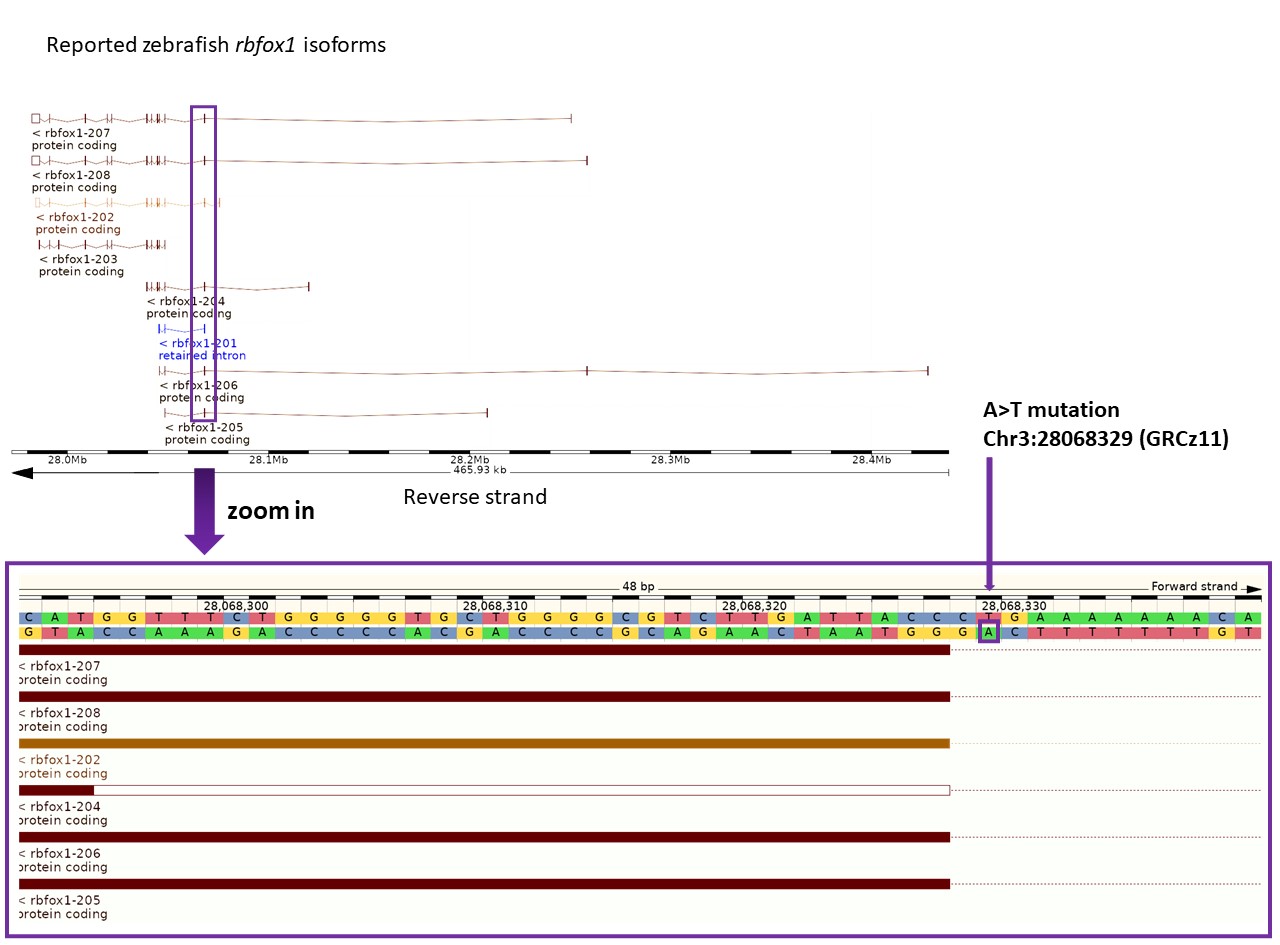

**Supplementary Figure 1. sa15940 mutation in *rbfox1* gene: effects in *rbfox1* expression in adult brain.** Top: *rbfox1* isoforms described in zebrafish (Ensembl database, GRCz11). Bottom: sa15940 is a point mutation (A>T, Chr3:28068329, GRCz11) situated in an intronic splicing region affecting all *rbfox1* protein-coding isoforms described in zebrafish except for *rbfox1*-203.

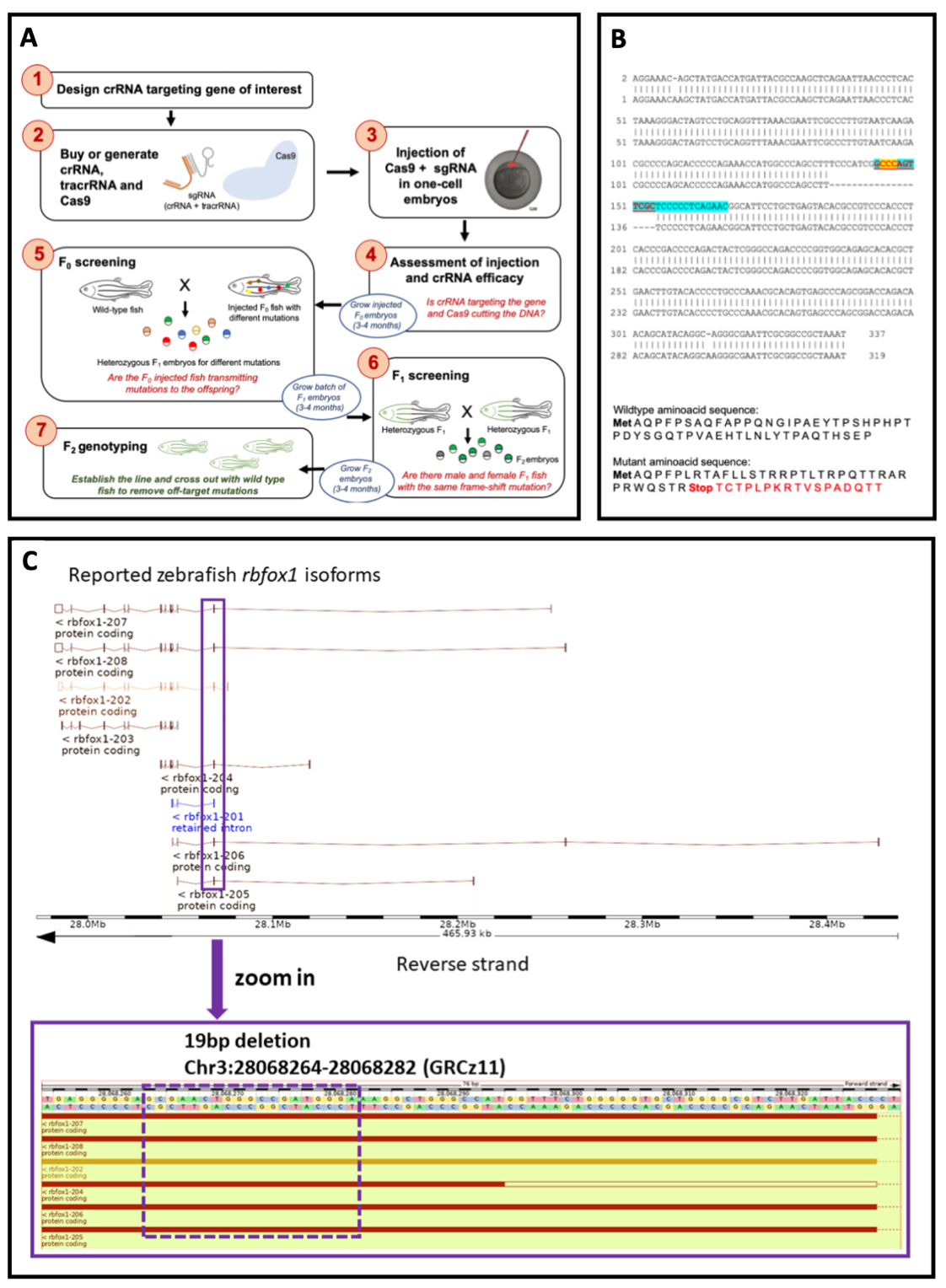

**Supplementary Figure 2**. **A) Procedure overview.** Generation of stable *rbfox1* loss of function zebrafish using CRISPR/Cas9 technology. **B)** **Confirmation of frameshift mutation for the generation of *rbfox1^del19^* loss-of-function line**. Top: Comparison of wild-type (top) and mutant (bottom) sequences for *rbfox1*. crRNA is highlighted in blue. PAM sequence is highlighted in yellow. The restriction site (that is disrupted in the F_0_ screening) appears in red, underlined font. Bottom: Comparison of wild-type (top) and mutant (bottom) amino acid sequences. Mutant sequence generates an early STOP codon. **C)** **del19 mutation in *rbfox1* gene created by CRISPR/Cas9.** Top: *rbfox1* isoforms described in zebrafish (Ensembl database, GRCz11). Bottom: del19 is a 19 bp deletion (Chr3:28068264-28068282, GRCz11) situated in an exon affecting all *rbfox1* protein-coding isoforms described in zebrafish except for *rbfox1*-203.

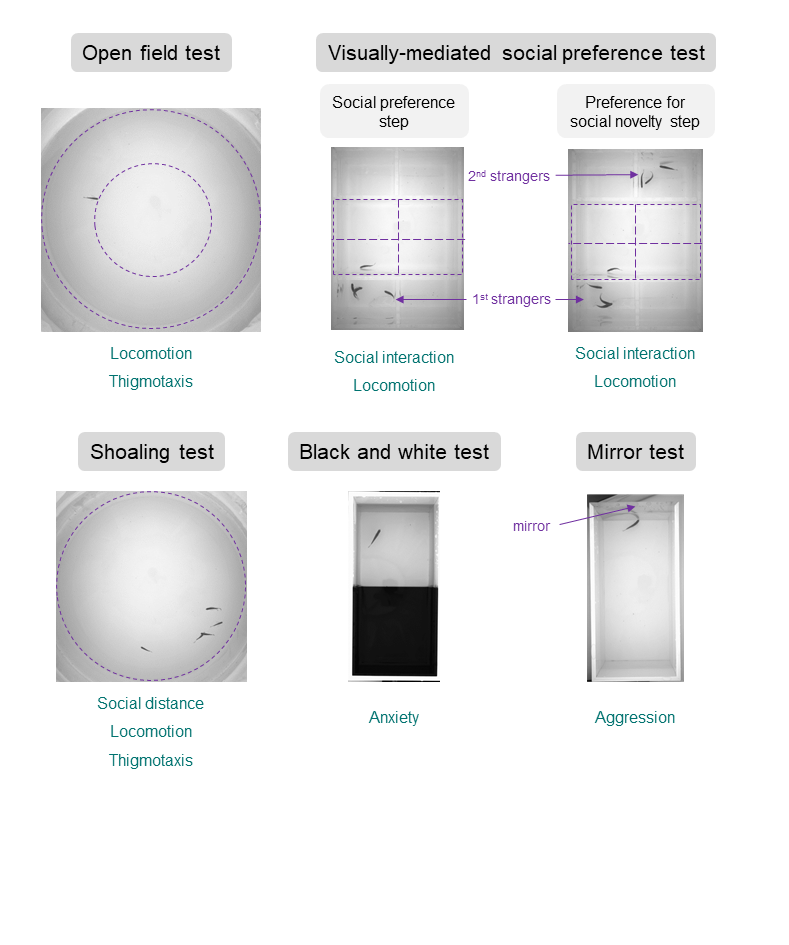

**Supplementary Figure 3**. **Battery of behavioural tests performed for both mutant lines.** The purple dotted lines indicate the areas where the tested fish was tracked. The type of behaviour that is assessed for each test is indicated in blue. Locomotion can assess hyperactivity, which is one component of psychiatric disorders like ADHD. Thigmotaxis is an anxiety-like behaviour and can be also considered a stereotypic behaviour associated with ASD. Social interaction and social distance: impairments in these tests can be related to ASD. Preference for the black side in the black and white test is another measure of anxiety-like behaviour. Aggression can be related to some types of aggressive behaviour in humans and co-occurs with many psychiatric disorders.

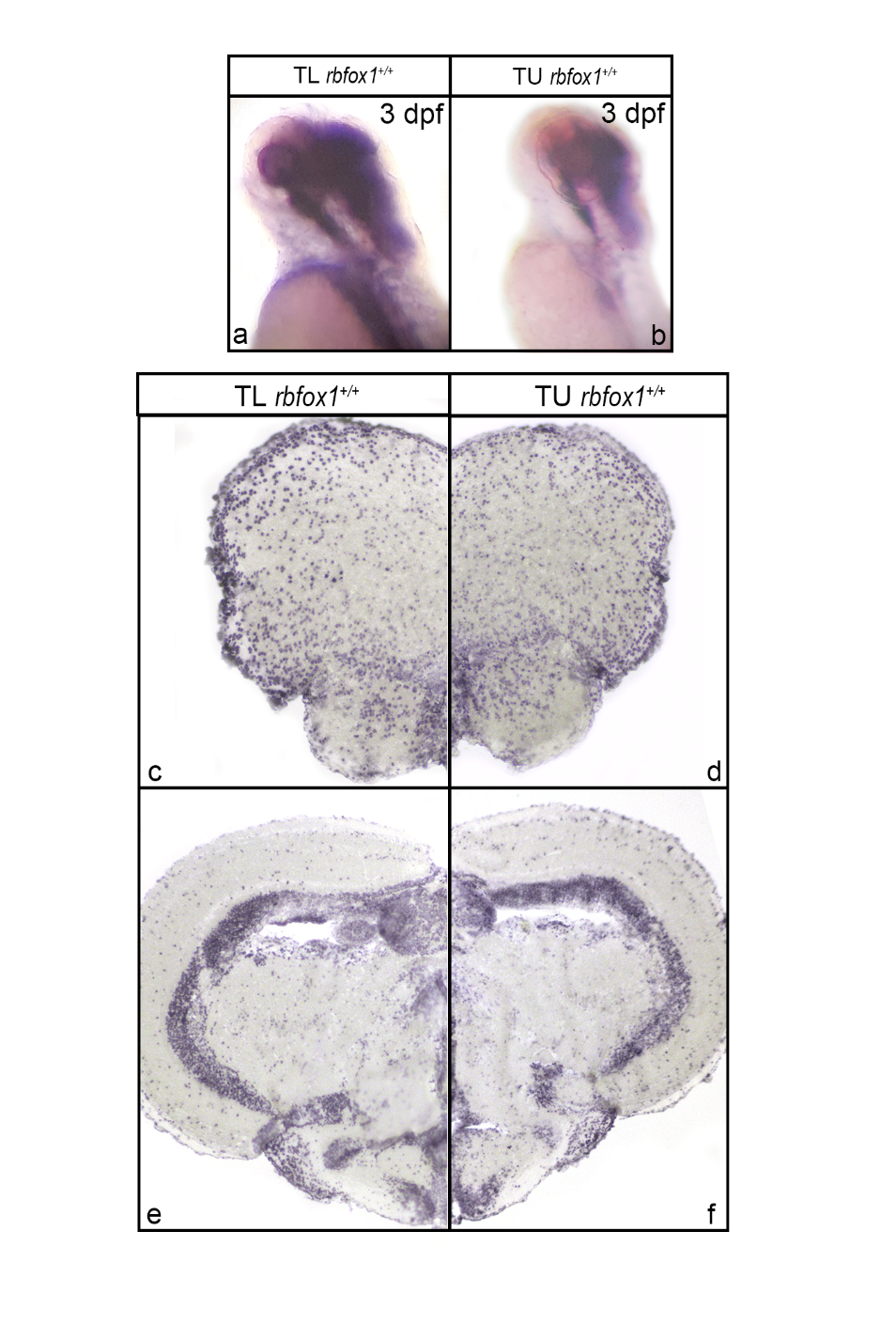

**Supplementary Figure 4**. **Comparison of r*bfox1* expression between TU *rbfox1^+/+^* and TL *rbfox1^+/+^*.** *rbfox1* whole-mount *in situ* hybridization on zebrafish larvae at 3 days post fertilization, (a) TL background, and (b) TU background. *rbfox1 in situ* hybridization on adult zebrafish brain, (c, e) TL background, and (d, f) TU background.

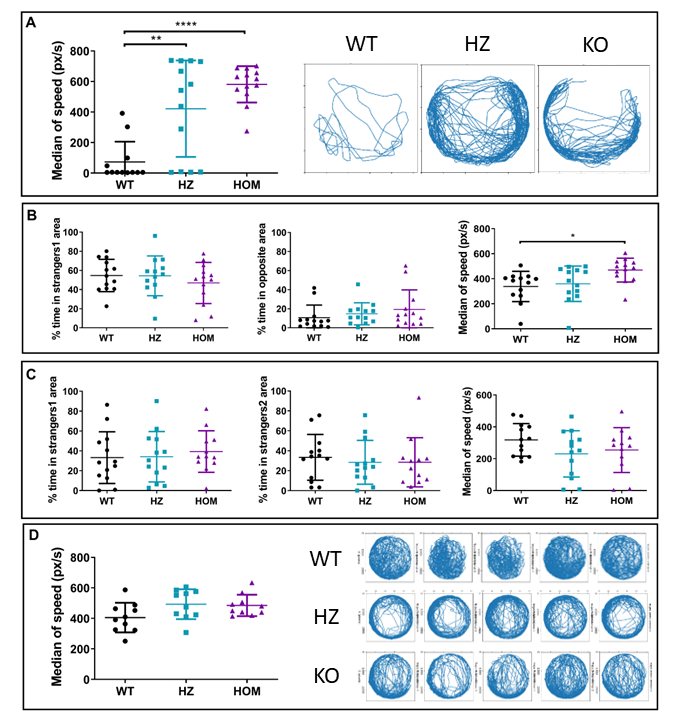

**Supplementary Figure 5**. **Behavioural alterations observed in *rbfox1*^sa15940^. A) Open field.** Left: median of speed during the open field test. One-way ANOVA followed by Tukey’s multiple comparison test. Right: representation of the trajectory of one individual of each genotype during the open field test. **B) Visually-mediated social preference test. Social preference step**. Time spent in the area close to the 1^st^ strangers and in the opposite area and median of speed during this step of the VMSP test. Kruskal-Wallis followed by Dunn’s multiple comparisons test. **C) Visually-mediated social preference test. Preference for social novelty step**. Time spent in the area close to the 1^st^ and to the 2^nd^ strangers and median of speed during this step of the VMSP test. Kruskal-Wallis followed by Dunn’s multiple comparisons test. **D) Shoaling test**. Left: median of speed during the shoaling test. Right: representation of the trajectory of five individual of each genotype during the open field test. n = 13 WT, 13 HZ and 13 HOM for all tests except for the shoaling test. For the shoaling test: n = 2 groups of 5 individuals per genotype. * p < 0.05; ** p < 0.01; **** p < 0.0001. Mean ± SD. HOM, *rbfox1*^sa15940/sa15940^ fish; HZ, *rbfox1* ^sa15940/+^ fish; WT, TL *rbfox1^+/+^*.

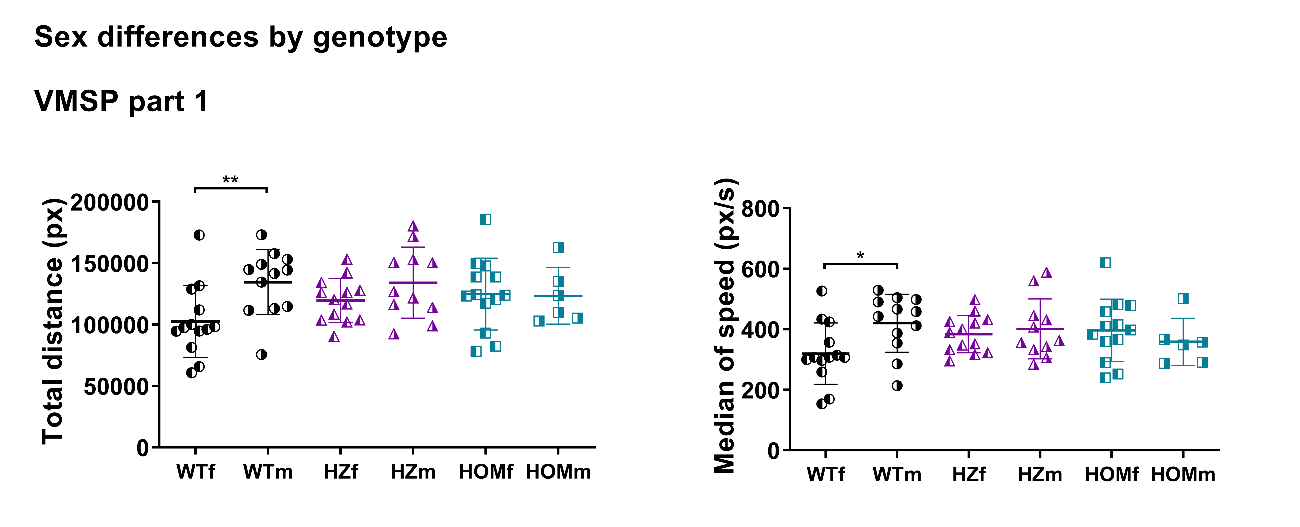

**Supplementary Figure 6. Significant behavioural alterations observed between sexes in the sex-separated *rbfox1*^sa15940^ batch.** Total distance travelled and median of speed during the first step of the visually-mediated social preference (VMSP) test. Kruskal-Wallis followed by Dunn’s multiple comparisons test. HOM, *rbfox1*^sa15940/sa15940^ fish; HZ, *rbfox1* ^sa15940/+^ fish; WT, wild-type TL. The *f* subindex indicates the female group and the *m* subindex indicates the male group. n = 13 WTf, 13 HZf, 13 HOMf, n = 12 WTm, 11 HZm, 9 HOMm.
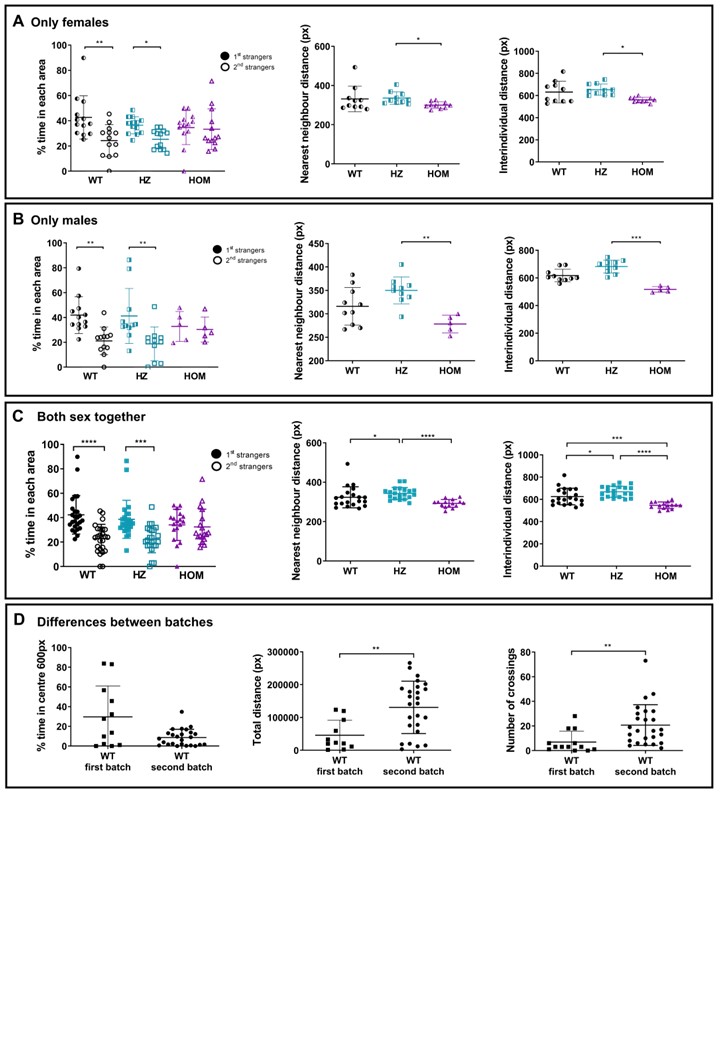

**Supplementary Figure 7. Behavioural alterations observed in the sex-separated *rbfox1*^sa15940^ batch considering all individuals in the analyses. A) Significant differences found between genotypes in the female batch of experiments performed with the *rbfox1*^sa15940^ line**. Left: Time spent in the areas close to the 1^st^ or 2^nd^ strangers during the preference for social novelty step of the VMSP test. n = 13 WT, 13 HZ, 13 HOM. Middle and right: mean of nearest neighbour and interindividual distance during the shoaling test. Kruskal-Wallis followed by Dunn’s multiple comparisons test. n = 10 WT, 10 HZ, 10 HOM. **B) Significant differences found between genotypes in the male batch of experiments performed with the *rbfox1*^sa15940^ line**. Left: Time spent in the areas close to the 1^st^ or 2^nd^ strangers during the preference for social novelty step of the VMSP test. n = 12 WT, 11 HZ, 9 HOM. Middle and right: mean of nearest neighbour and interindividual distance during the shoaling test. Kruskal-Wallis followed by Dunn’s multiple comparisons test. n = 10 WT, 10 HZ, 5 HOM. **C) Significant differences found between genotypes in the sex-separated batch of experiments performed with the *rbfox1*^sa15940^ line**. Left: Time spent in the areas close to the 1^st^ or 2^nd^ strangers during the preference for social novelty step of the VMSP test. n = 25 WT, 24 HZ, 18 HOM. Middle and right: mean of nearest neighbour and interindividual distance during the shoaling test. Kruskal-Wallis followed by Dunn’s multiple comparisons test. n = 20 WT, 20 HZ, 15 HOM. **D) Comparison of WT behaviour between the first and the second batch of experiments.** Left and middle: time spent in the centre of the arena and total distance travelled during the open field test. Right: number of crossings during the black and white test. Mann-Whitney U test. Mean ± SD. * p < 0.05; ** p < 0.01, *** p < 0.001, **** p < 0.0001.

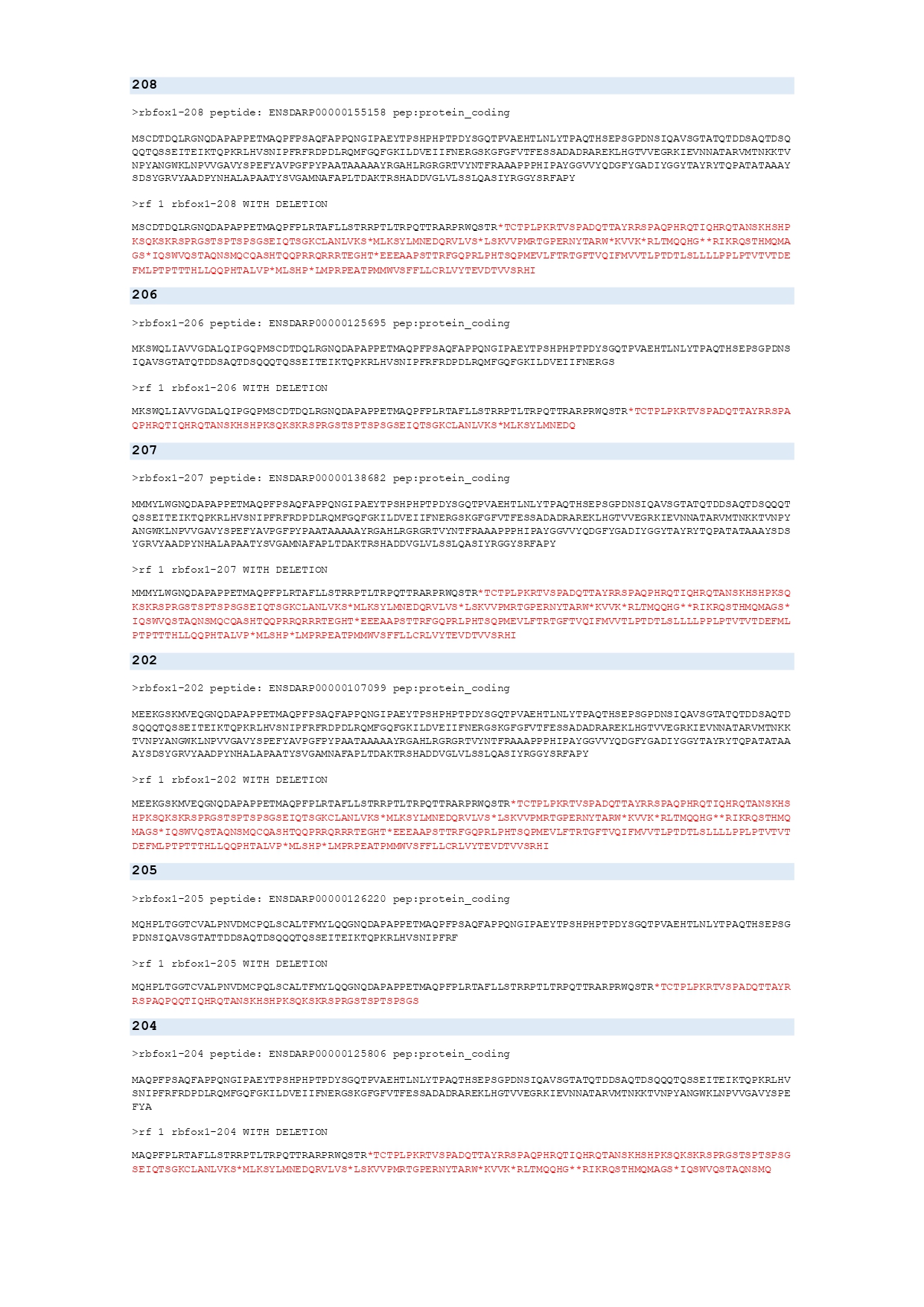

**Supplementary Figure 8**. Predicted protein sequence of *rbfox1* isoforms with and without the exonic frameshift caused by 19 bp deletion. Isoforms are named according to Ensembl nomenclature and the protein sequences were obtained from the Ensembl database (GRCz11, <https://www.ensembl.org/Danio_rerio>). For each isoform, the predicted effect of the 19 bp deletion on the protein sequence has been described. In black: protein sequence. In red: non-translated amino acids due to the premature presence of a stop codon.

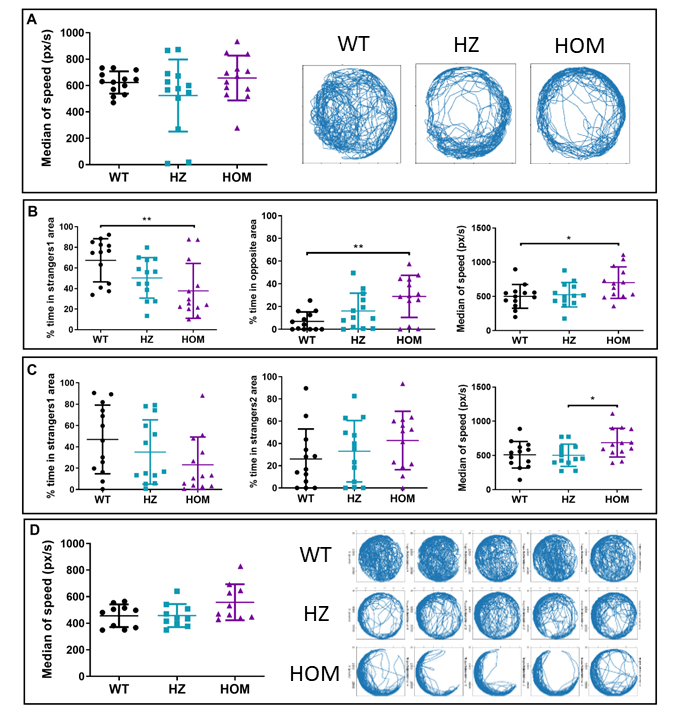

**Supplementary Figure 9**. **Behavioural alterations observed in *rbfox1*^del19^.** **A) Open field.** Left: median of speed during the open field test. One-way ANOVA followed by Tukey’s multiple comparison test. Right: representation of the trajectory of one individual of each genotype during the open field test. **B) Visually-mediated social preference test. Social preference step**. Time spent in the area close to the 1^st^ strangers and in the opposite area and median of speed during this step of the VMSP test. Kruskal-Wallis followed by Dunn’s multiple comparisons test. **C) Visually-mediated social preference test. Preference for social novelty step**. Time spent in the area close to the 1^st^ and to the 2^nd^ strangers and median of speed during this step of the VMSP test. Kruskal-Wallis followed by Dunn’s multiple comparisons test. **D) Shoaling test**. Left: median of speed during the shoaling test. Right: representation of the trajectory of five individual of each genotype during the open field test. n = 13 WT, 13 HZ and 13 HOM for all tests except for the shoaling test. For the shoaling test: n = 2 groups of 5 individuals per genotype. * p < 0.05; ** p < 0.01. Mean ± SD. HOM, *rbfox1*^del19/ del19^ fish; HZ, *rbfox1*^del19/+^ fish; WT, TU *rbfox1^+/+^*.

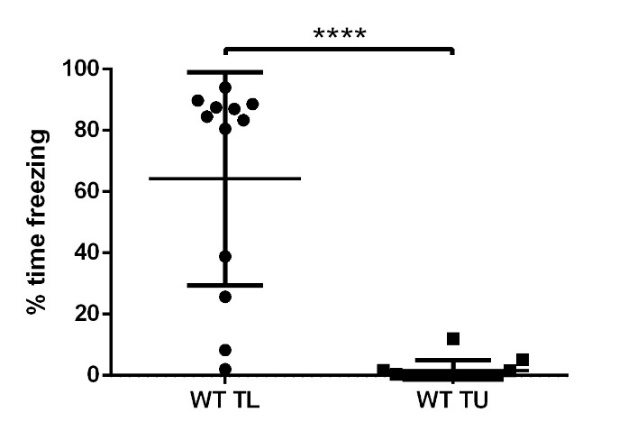

**Supplementary Figure 10. Comparison of the time freezing during the open-field test between the two wild-type lines used as controls in the behavioural experiments**. WT, wild-type; TL, Tübingen Long-fin line; TU, Tübingen line. **** p < 0.0001, Mann-Whitney U test. Mean ± SD.

3. **SUPPLEMENTARY TABLES**

**Supplementary Table 1. Sequence of the primers used for the RT-qPCRs.**

| **Gene** | **Sequence of forward primer** | **Sequence of reverse primer** | **Length of the product** |
| --- | --- | --- | --- |
| *rpl13a* | 5′-Tctggaggactgtaagaggtatgc-3’ | 5′-AGACGCACAATCTTGAGAGCAG-3’ | 148 |
| *eef1a1a* | 5′-GAGGAAATCACCAAGGAAGTCAG-3’ | 5′-TTGAACCAGCCCATGTTTGAG-3’ | 134 |
| *ube2a* | 5′-TGACTGTTGACCCACCTTACAG-3’ | 5′-CAAATAAAAGCAAGTAACCCCG-3’ | 265 |
| *rbfox1** | 5′-AACAGCATACAGGCGGTCTC-3’ | 5′-GGCTGCGTTTTGATTTCTGT-3’ | 110 |
| *rbfox1l* | 5′-TCCCAGTTTACCCTCAGACG-3’ | 5′-AGGTGGAGGATATGAAGGGG-3’ | 134 |
| *rbfox2* | 5′-CCGTATGCTAACGGGGACAG-3’ | 5′-ACCCTGGAACTGCGTAGAAC-3’ | 106 |
| *rbfox3a* | 5′-CCCTCACCATCTACACACCA-3’ | 5′-TCCGAAGAGTCTGATGGCTG-3’ | 160 |
| *rbfox3b* | 5′-GAGTCCACAGAAAAGCAGCA-3’ | 5′-TCTCCACGTCTAAGATCTTCCC-3’ | 118 |

* These *rbfox1* primers amplify all the described isoforms in GRCz11 except for *rbfox1*-203

**Supplementary Table 2. crRNA + tracrRNA for generation of *rbfox1^del19^* mutants**. Genomic regions targeted by the crRNA (black) and PAM sequence (red) with the restriction sites (underlined).

| **Gene name exon #** | **Restriction enzyme** | **DNA region targeted**  **by crRNA and PAM** | **crRNA + tracrRNA overlapping sequence (in blue)** |
| --- | --- | --- | --- |
| *rbfox1*  exon 2 | MwoI | GCCCAGTTCGCTCCCCCTCAGAAC | GUUCUGAGGGGGAGCGAACUGUU UUAGAGCUAUGCUGUUUUG |

4. **REFERENCES**

1	Wullimann MF, Rupp B, Reichert H. Neuroanatomy of the Zebrafish Brain. *Neuroanat Zebrafish Brain* 1996. doi:10.1007/978-3-0348-8979-7.

2	Romero-Ferrero F, Bergomi MG, Hinz RC, Heras FJH, de Polavieja GG. idtracker.ai: tracking all individuals in small or large collectives of unmarked animals. *Nat Methods 2019 162* 2019; **16**: 179–182.

3	Parker MO, Brock AJ, Millington ME, Brennan CH. Behavioural phenotyping of casper mutant and 1-Pheny-2-Thiourea treated adult zebrafish. *Zebrafish* 2013; **10**: 466–471.

4	Carreño Gutiérrez H, Colanesi S, Cooper B, Reichmann F, Young AMJ, Kelsh RN *et al.* Endothelin neurotransmitter signalling controls zebrafish social behaviour. *Sci Rep* 2019; **9**. doi:10.1038/s41598-019-39907-7.

5	Ruhl N, McRobert SP, Currie WJS. Shoaling preferences and the effects of sex ratio on spawning and aggression in small laboratory populations of zebrafish (Danio rerio). *Lab Anim (NY)* 2009; **38**: 264–269.

6	Norton WHJ, Stumpenhorst K, Faus-Kessler T, Folchert A, Rohner N, Harris MP *et al.* Modulation of fgfr1a signaling in zebrafish reveals a genetic basis for the aggression-boldness syndrome. *J Neurosci* 2011; **31**: 13796–13807.
